# High-resolution integrative analysis allows characterization and spatial annotation of normal and cancer-associated colon fibroblasts

**DOI:** 10.1101/2025.07.29.667377

**Authors:** Emma W Viitala, Ella Salminen, Mitro Miihkinen, Niki Chalkidi, Athanasia Stavropoulou, Toni T Lemmetyinen, Pekka Päivinen, Tuomas Kaprio, Jaana Hagström, Caj Haglund, Tomi P. Mäkelä, Tero Aittokallio, Pekka Katajisto, Vasiliki Koliaraki, Saara Ollila

**Author notes:** Corresponding authors: Emma Viitala,; Saara Ollila.

## Abstract

Fibroblasts represent key regulators of colon homeostasis, and cancer-associated fibroblasts (CAFs) play pivotal roles in colorectal cancer (CRC). Despite their critical influence, a consensus view of adult human colon fibroblast and CRC CAF heterogeneity, spatial organization, and developmental trajectories is currently lacking. Here, we address this gap by performing a comprehensive characterization of colonic fibroblasts and CRC CAFs. Using large-scale integration of single-cell RNA-sequencing datasets from normal colon and CRC we mapped the fibroblast subpopulations. Spatial transcriptomics, immunohistochemistry, and *in situ* hybridization were used for validation and computational analyses to predict developmental trajectories and transcription factors underlying CAF activation. Subepithelial myofibroblasts (SEMFs), mucosa-associated fibroblasts (MAFs), and submucosa-associated fibroblasts (SAFs) were identified as the main colon fibroblast subtypes, and in mice, further divided into location-based subclusters. We also identified a novel colon fibroblast subset: muscle-embedded interstitial fibroblasts (MIFs). CRCs contained normal fibroblasts as well as four cancer-specific CAF populations: inflammatory CAFs (iCAFs), matrix CAFs (mCAFs), and two precursor CAF (preCAF) subtypes. Our data suggested that iCAFs originate from SEMFs through a preCAF1 intermediate phenotype, while mCAFs derive from SAFs/MIFs via preCAF2s. Transcription factors PRRX1, MAFB, and TWIST1 were uncovered as potential regulators of CAF identity and CTHRC1 was identified as a specific and sensitive pan-CAF marker in CRC. Our study presents a detailed framework for understanding colon fibroblast and CRC CAF heterogeneity. This work lays the groundwork for future research into the roles and potential therapeutic relevance of different CRC CAF subsets.

## INTRODUCTION

Recent advances in single-cell technologies have revealed a vast transcriptional and functional heterogeneity of fibroblasts across various tissues [1–3], placing fibroblasts as a subject of intense research. In the colon, fibroblasts represent key regulators of epithelial turnover, and three main fibroblast subtypes have been characterized mostly in studies investigating mouse tissues: (1) *Pdgfra*^high^ fibroblasts, promoting epithelial cell differentiation e.g. through expression of BMP ligands, and *Pdgfra*^low^ fibroblasts, which further divide into (2) *Pi16*-*Cd81*- and (3) *Pi16*+ *Cd81*+ subsets, supporting stem cell function by secreting BMP inhibitors and WNT agonists [4–11]. Variable nomenclature has been proposed for the identified colon fibroblast subsets, based on factors such as cluster numbers, markers, topology, morphology, or functional properties (S.Table 3). For example, *Pdgfra*^high^ cells have been referred to as subepithelial myofibroblasts (SEMFs) or telocytes; *Pdgfra*^low^ *Pi16*-*Cd81*-fibroblasts as mucosa-associated fibroblasts (MAFs) or crypt bottom fibroblast 1 (CBF1); and *Pdgfra*^low^ *Pi16*+ *Cd81+* cells as trophocytes or crypt bottom fibroblast 2 (CBF2) [5, 6, 12]. Human colon fibroblasts appear to exhibit transcriptional similarities to mouse cells [4, 13, 14], but a systematic analysis and annotation of adult human colon fibroblast subsets aligned with the current understanding of the mouse fibroblast subsets is lacking.

In addition to maintaining homeostatic tissues, fibroblasts play key roles in carcinogenesis, thus being increasingly recognized as potential drug targets in cancer [15, 16]. Cancer-associated fibroblasts (CAFs) are known to regulate tumor growth, immune responses, and drug resistance [17–19], but are heterogeneous and exhibit both tumor-promoting and tumor-restraining functions [20–22], emphasizing the acute need for careful characterization and annotation of different CAF subtypes. In colorectal cancer (CRC), the third most common and second most fatal cancer type worldwide (Cancer today 2022), the CAF-rich, mesenchymal CRC consensus molecular subtype 4 (CMS4) [23] is associated with poor survival and resistance to chemotherapy [24, 25], highlighting the importance of understanding the functional heterogeneity of CRC CAFs.

Several studies addressing the CAF heterogeneity in CRC have been conducted, reporting several CRC CAF subtypes, including inflammatory (iCAF), myofibroblastic/matrix (mCAFs), antigen-presenting (apCAFs), vascular (vCAFs), chemo-CAFs, tumor-like (tCAFs), and proliferative (pCAFs), along with tens of other, marker-based names (S.Table 4). However, a consensus view of CRC CAF subsets and their prognostic significance is currently lacking. Furthermore, how normal tissue-resident colon fibroblast subsets relate to the developmental trajectories of CRC CAFs is currently unknown. One major bottleneck preventing reproducible characterization of CRC CAF subsets is the lack of consistent annotation of tissue-resident colon fibroblasts and other mesenchymal cells, such as pericytes, smooth muscle cells (SMCs), and myofibroblasts, leading to all mesenchymal cells in resected cancers often being labelled as CAFs [2, 3, 26–35]. This, together with limitations such as small sample size and/or low resolution [36–42], has made it difficult to create a coherent view on CRC CAF subsets. In spatial analyses, low resolution and lack of specific markers have challenged the formation of a clear view on CAF subtype localization [3, 32, 42, 43]. Finally, despite providing important information about the general properties of CAFs, cross-tissue atlases fail to capture the tissue-specific characteristics of CAF subtypes [2, 3, 37, 40, 42]. Therefore, a benchmarking analysis of CRC CAF subsets is urgently needed.

In this study, we adopt an integrative approach to fill the gaps in the normal colon fibroblast and CRC CAF characterization. First, we generate a high-resolution map of mouse colon fibroblast subclusters. Second, using the mouse analysis as a benchmark, we annotate human colon fibroblasts and validate their localization with spatial transcriptomics. Finally, we annotate CRC CAF subsets, establish their potential differentiation trajectories, and identify potential regulators underlying CAF development. Our work provides a solid foundation for future research into colon fibroblast heterogeneity and CAF subset-specific investigations in CRC.

## RESULTS

### Analysis and comparison of mouse and human colon fibroblast subsets

To investigate the heterogeneity of colonic fibroblasts, we first performed single-cell RNA-sequencing (scRNA-seq) of mouse colonic mesenchyme and analyzed the fibroblasts and smooth muscle cells (SMCs). Consistent with previous analyses [4, 5, 7, 8], the fibroblasts clustered into three main subsets: (1) *Pdgfra*^high^, (2) *Pdgfra*^low^ *Pi16*-*Cd81*-, and (3) *Pdgfra*^low^ *Pi16*+ *Cd81*+ populations (S1A and S1B Fig). Interestingly, however, the *Pdgfra*^low^ *Pi16*-*Cd81*-cluster was further divided into two subsets (S1A and S1C Fig), indicating heterogeneity within the main colon fibroblasts subtypes, as proposed previously [4, 11]. To investigate this with higher resolution, we integrated our data with four previously published datasets containing mouse colon mesenchymal cells [4, 5, 7, 8] (S1D Fig). The integration resulted in 36.085 cells including fibroblasts (n = 25.937), SMCs, pericytes, endothelial cells, glial cells, and epithelial cells (Fig 1A and 1B, S2A and S2B Fig). Using this approach, mouse colonic fibroblasts clustered into seven sub-populations, including three *Pdgfra*^high^ and four *Pdgfra*^low^ subsets (Fig 1B, S2A Fig). Among the *Pdgfra*^high^ fibroblasts, we observed two clusters, a *Lama5*+ and a *Wif1*+ cluster, both expressing *Foxl1, F3, Wnt5a, Bmp5,* and *Bmp7*, suggesting that these clusters represent the SEMF/telocyte population [5, 6, 12]. The third *Pdgfra*^high^ cluster expressed *Kcnn3*+ but did not match the expression profile of previously identified colon fibroblasts or other mesenchymal cells (Fig 1B, S2B Fig). The *Pdgfra*^low^ cells consisted of three *Pi16*-*Cd81*-clusters, representing the previously identified mucosa-associated fibroblasts (MAFs) [8], and one *Pi16*+ *Cd81*+ population (Fig 1B). The mucosal WNT-BMP gradient (Fig 1C) confirmed that the observed fibroblast populations (except for the *Pdgfra*^high^ *Kcnn3*+-cluster) represented the *Pdgfra*^low^, stemness-supporting fibroblasts, and the *Pdgfra*^high^, differentiation-promoting fibroblasts, previously shown to be distributed along the basal-luminal axis of colonic mucosa [5, 6, 12].

**Fig 1.**
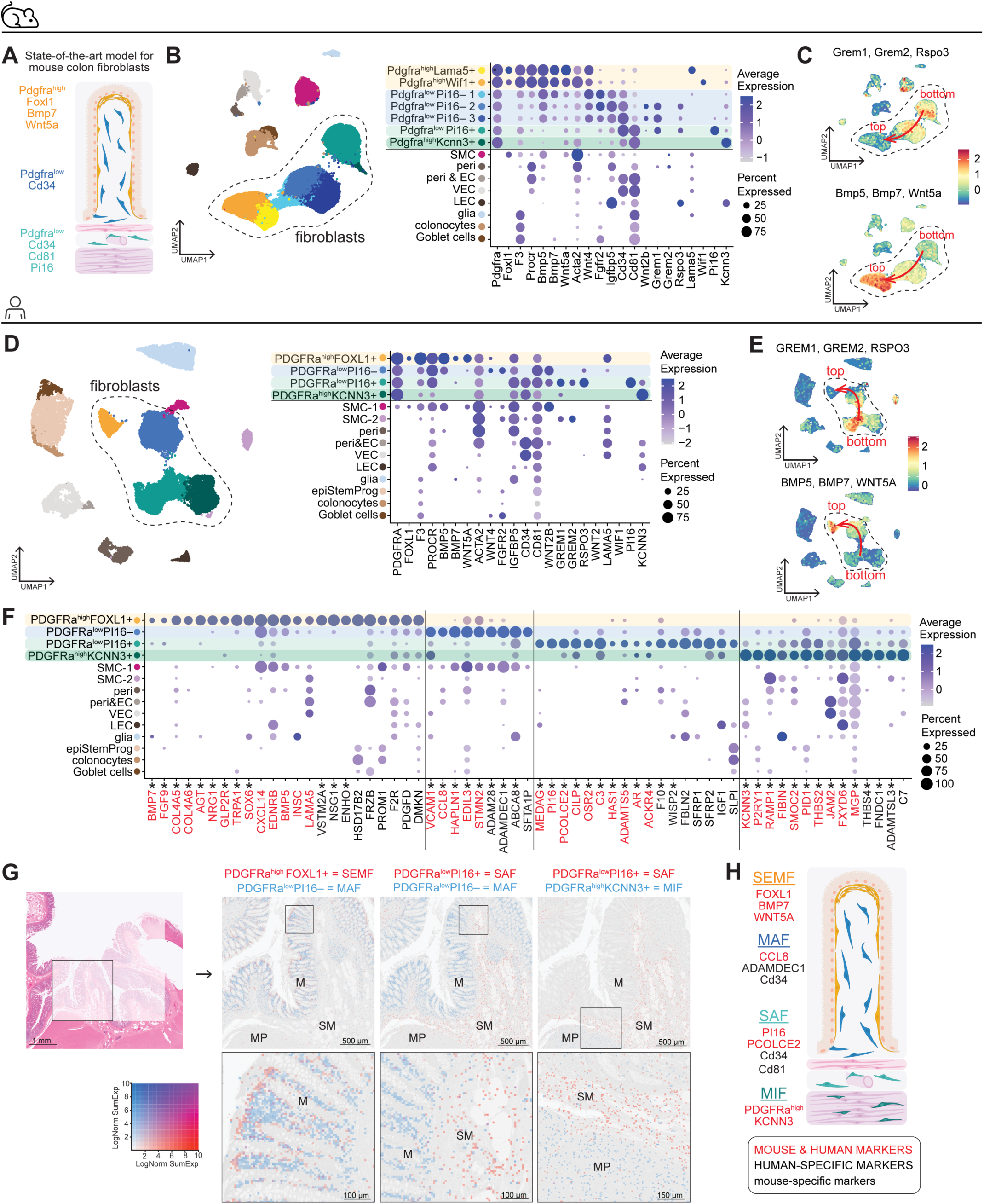
Annotation of mouse and human colon fibroblast subtypes using high-throughput analysis of scRNA-seq data. (A) Current model of mouse colon fibroblast subsets. (B) Uniform manifold approximation and projection (UMAP) plot of integrated mouse colon scRNA-seq data and (left) dot plot depicting markers of identified colonic fibroblast subpopulations in mouse (right). See also S2B Fig. (C) Signature feature plots of indicated WNT and BMP genes. (D) UMAP of integrated human colon scRNA-seq data (left) and dot plot of the corresponding human markers shown for mice in B (right). See also S2D Fig. (E) Signature feature plots of indicated WNT and BMP genes. (F) Dot plot depicting top marker genes for each identified human colon fibroblast subpopulation. Common markers between mouse and human are marked in red. Asterisk (*) indicates genes included in the signatures used in G. See also S2 Table. (G) Signatures of human fibroblast subpopulations (marked by asterisk in F) projected onto spatial transcriptomics data of normal human colon. M, mucosa; SM, submucosa: MP, muscularis propria. (H) Updated model of mouse and human colon fibroblast subsets. SEMF, subepithelial myofibroblast; MAF, mucosa-associated fibroblasts; SAF, submucosa-associated fibroblasts; MIF, muscle-embedded interstitial fibroblasts; SMC, smooth muscle cells; peri, pericytes; EC, endothelial cells; VEC, vascular endothelial cells; LEC, lymphatic endothelial cells; epiStemProg, epithelial stem and progenitor cells.

To address how human colon fibroblasts correspond to the mouse counterparts, we analyzed scRNA-seq data of human colonic normal mucosal cells from four datasets (CRC16, KUL3, KUL5, SMC) sampled during surgery from CRC patients (n = 36) (S1E Fig) [26, 44]. Notably, the normal mesenchymal cells from these datasets have not been previously examined in detail while some studies have classified all (including normal colon-derived) fibroblasts from these datasets as CAFs [33, 38, 39], calling for clarification in annotation. After filtering out the immune cells (to match the mouse analysis), the integration resulted in 22.215 cells, including fibroblasts (n = 11.702), SMCs, pericytes, endothelial cells, glial cells, and epithelial cells, with similar contributions across datasets (Fig 1D, S2C and S2D Fig). Of note, also *KIT*+ intestinal cells of Cajal (ICCs) [45] and *CCL19*+ reticular fibroblasts [46] were identified, but not in sufficiently high numbers to form separate clusters (S2E Fig).

The human fibroblasts divided into two *PDGFRA*^high^ clusters, one expressing *FOXL1*+ and the other *KCNN3*+, and two *PDGFRA*^low^ populations that differed by *PI16* expression (Fig 1D). Except for the *KCNN3*+ cells, the BMP-WNT gradient suggested a basal-luminal distribution, as in the mouse analysis (Fig 1E). Despite some species-specific differences for individual marker genes (S2F and S2G Fig), the overall expression patterns allowed correlation of mouse and human fibroblast subsets (S2H Fig, Fig 1F). The human *PDGFRA*^high^ *FOXL1*+ cluster expressed *BMP5* and *BMP7*, consistently with SEMF/telocyte identity shared with mouse cells. The human *PDGFRA*^low^ *PI16*-cells also expressed several marker genes shared with their mouse counterparts, including e.g., *CCL8,* a cytokine previously reported as a colon fibroblast subset-specific marker [4]. Unlike in mice, the human *PDGFRA*^low^ cells did not show differential expression of *CD81*, but expression of markers such as *ACKR4, C3,* and *PCOLCE2* suggested shared identity of *PDGFRA*^low^ *PI16*+ cells with the mouse *Pdgfra*^low^ *Pi16*+ *Cd81*+ cells (Fig 1F, S2H Fig). Interestingly, in mice, expression of *Grem1* and *Rspo3* was noted both in the *Pdgfra*^low^ *Pi16*-*Cd81*- and the *Pdgfra*^low^ *Pi16*+ *Cd81*+ cells (Fig 1B), while in humans *GREM1* and *RSPO3* were primarily derived from *PDGFRA*^low^ *PI16*+ cells, and the *PDGFRA*^low^ *PI16*-cells instead expressed the canonical Wnt ligand *WNT2B* (Fig 1d). Thus, the exact molecular mechanisms promoting canonical Wnt signaling in the colonic stem cell niche may slightly differ in mice and humans. Altogether, the gene signatures and growth factor expression patterns of human fibroblast subclusters suggested a similar organization to the mouse colon.

Next, we set out to annotate the identified human colon fibroblast clusters according to their spatial distribution by mapping the cluster-specific expression signatures (Fig 1F and 1G) in a high-resolution Visium HD spatial transcriptomics dataset of human colon [47]. SEMFs have been described as fibroblasts with expression of myofibroblast markers and a subepithelial location throughout the gastrointestinal tract, including the small intestine and colon [48]. Consistently, the *PDGFRA*^high^ *FOXL1*+ cells were enriched directly beneath the epithelial cells, especially in the luminal side of colonic mucosa, and expressed the myofibroblast marker *ACTG2* (S2D Fig), confirming the SEMF identity (Fig 1G). The *PDGFRA*^low^ *PI16*-cells localized within the mucosal lamina propria, and we named these cells mucosa-associated fibroblasts (MAFs). The *PDGFRA*^low^ *PI16*+ cluster genes were highly enriched in the submucosa, consistent with previous analysis of the mouse *Pdgfra*^low^ *Pi16*+ *Cd81*+ cells [8], and we named these cells submucosa-associated fibroblasts (SAFs). Interestingly, the *PDGFRA*^high^*KCNN3*+ fibroblasts were embedded among the SMCs in the muscularis propria, and this cluster was named muscle-embedded interstitial fibroblasts (MIFs). Thus, we propose a coherent, location-based nomenclature for human and mouse fibroblast subsets, validated by spatial transcriptomics in human tissue and supported by previous literature and shared markers in mice (S3 Table, Fig 1H).

### Sub-clustering of MAFs and SEMFs along the crypt-apex and proximal-distal axes in mouse colon

Next, we investigated the sub-clusters of MAFs identified in the mouse colon. Three separate clusters, MAF-1, MAF-2 and MAF-3, were identified (Fig 1B, Fig 2A). *Wnt4*, expressed by fibroblasts residing in the top (apex) part of the colonic lamina propria [9], was seen in the MAF-1 cluster, while MAF-2 and MAF-3 expressed *Grem1*, a known niche factor for stem cells (Fig 2B) [6, 10, 49–52]. Therefore, the MAF-1 cluster likely represents MAFs localized at the crypt top (Top MAF), whereas MAF-2 and MAF-3 reside at the bottom part of the mucosa, closer to the stem cell zone (Bottom MAF). Interestingly, one of the analyzed datasets contained annotated cells deriving from either proximal or distal parts of the colon [8]. In this data, the MAF-2 cluster was primarily derived from proximal colon samples and expressed *Tnc,* while MAF-3 cells originated from the distal colon samples and expressed high levels of *Cd55* (Fig 2C and 2D). Tissue staining confirmed Tenascin C (encoded by *Tnc*) expression throughout the top part of the mucosal stroma as previously shown [53], while the crypt bottom stroma only expressed Tenascin C in the proximal colon (Fig 2E). To investigate the similarity between colonic and small intestinal (SI) mesenchymal cells, we also performed scRNA-seq on mouse SI mesenchyme and analyzed the fibroblasts and SMCs (S3A and S3B Fig). We identified *Pdgfra*^high^ *Foxl1*+ SEMFs (S3C Fig), two *Pdgfra*^low^ *Pi16*-MAF clusters (S3D Fig), and *Pdgfra*^low^ *Pi16+ Cd81+* SAFs (S3E Fig). Consistent with the colon analysis, *Wnt4* expression separated Top MAFs from Bottom MAFs (S3D Fig). However, Bottom MAFs did not separate further, and *Tnc* and *Cd55* expressions were uniform across MAFs.

**Fig 2.**
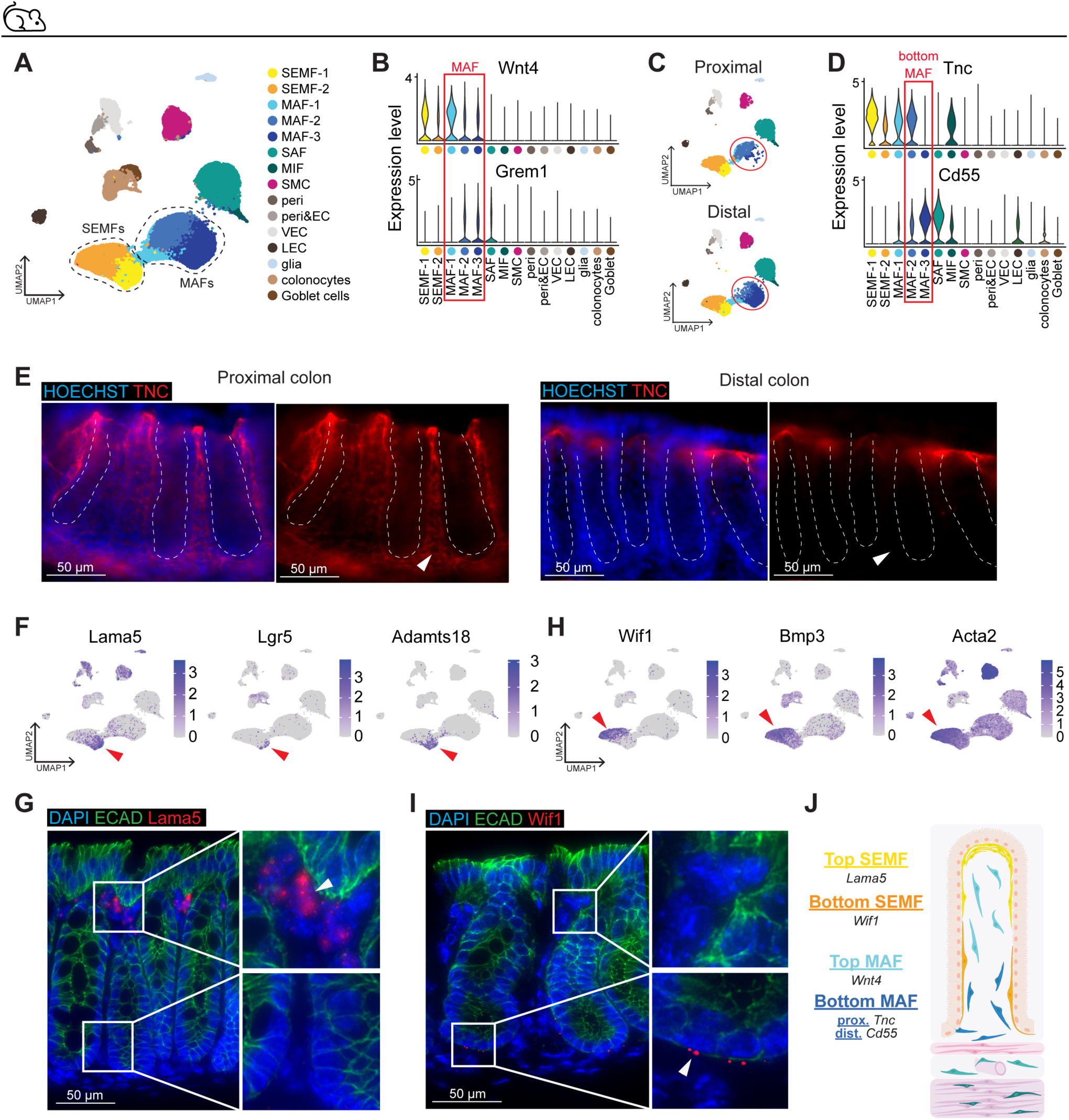
Sub-clustering mouse MAFs and SEMFs along the crypt-apex and proximal-distal axes. (A) UMAP of integrated mouse colon mesenchyme scRNA-seq data. SEMFs (subepithelial myofibroblasts) and MAFs (mucosa-associated fibroblasts) highlighted by dashed lines. (B) Violin plot of *Wnt4* and *Grem1* expression across the mesenchymal cell clusters. (C) UMAP of mouse colon mesenchymal cells from proximal and distal colons, MAFs highlighted by red circles. (D) Violin plots depicting differential expression of *Tnc* and *Cd55* in proximal (MAF-2) and distal (MAF-3) bottom MAFs. (E) Immunofluorescent staining of TNC in proximal and distal mouse colon. HOECHST: nuclei. (F) Feature plots of *Lama5, Lgr5* and *Adamts18* expression in mouse colon mesenchyme. Arrowheads point to Top SEMFs. (G) mRNA *in situ* hybridization of *Lama5* in mouse colon. (H) Feature plots of *Wif1*, *Bmp3,* and *Acta2* expression in mouse colon mesenchyme. Arrowheads point to Bottom SEMFs. (I) mRNA *in situ* hybridization of *Wif1* in mouse colon. (G and I), ECAD: E-cadherin, epithelial cells, DAPI: nuclei. (J) Updated model of mouse colon fibroblast subsets including MAF and SEMF subclusters and their markers.

Next, we addressed the heterogeneity of mouse colon SEMFs. SEMF-1 and SEMF-2 populations were identified, corresponding to *Lama5*+ and *Wif1*+ SEMFs, respectively (Fig 1B; Fig 2A, 2F and 2H). The *Lama5*+ SEMFs expressed *Wnt4,* indicating crypt top identity (Fig 2B) [9]. Consistently, the *Lama5*+ SEMF markers *Lgr5* and *Adamts18* (Fig 2F) were previously shown to be expressed in villus tip fibroblasts in the mouse small intestine [54]. Indeed, *Lama5* mRNA was localized at the top part of the colonic lamina propria (Fig 2G), and the *Lama5*+ SEMFs were annotated as “Top SEMFs”. The *Wif1*+ SEMF population, on the other hand, expressed *Bmp3* and high levels of *Acta2* (Fig 2H). Since *Bmp3* encodes for a BMP inhibitor [55] and *Pdgfra*^high^ *Acta2*^high^ cells have previously been shown to be located in the bottom colonic stroma [11], we hypothesized that these cells could localize at the stem cell zone. Indeed, *Wif1* mRNA was localized to stromal cells adjacent to the crypt bottom (Fig 2I), and we named these cells “Bottom SEMFs”. In SI, we only observed one SEMF population (S3C Fig), and *Wnt4* was expressed uniformly throughout the cluster (S3D Fig). However, the Top SEMF markers *Lama5* and *Adamts18* were enriched on one edge of the SEMF cluster, while *Wif1* was enriched on the other edge (S3C Fig), suggesting a similar bottom-to-top gradient in the SI SEMFs that could be more clearly separated in a larger dataset.

In summary, integration of multiple datasets allowed characterization of fibroblast subtypes of the mouse colon with high resolution, revealing differences along the basal-luminal and proximal-distal axes (Fig 2J). Interestingly, however, these subclusters were less clear in the human analysis, indicating subtle species-specific differences or lower resolution due to a smaller sample size (S4 Fig).

### SMC subtypes in mouse and human colon

A thin layer of smooth muscle, the muscularis mucosae (MM), separates colonic mucosa from the submucosa, surrounded by the circular and longitudinal smooth muscle layers constituting the muscularis propria (MP). In literature, the colonic stromal cells expressing SMC markers such as *ACTA2, MYH11, ACTG2,* and *DES* are variably referred to as SMCs, myofibroblasts, or myocytes [4, 5, 9, 10, 13, 56, 57], calling for clarity in annotation. We first subsetted and re-clustered the SMCs from the integrated mouse dataset, resulting in two clusters expressing SMC markers but differing in expression of *Hhip* (n = 3.636) (Fig 3A and 3B). The *Hhip* negative (*Hhip*-) cluster expressed *Grem1*, *Grem2*, *Rspo3,* and *Chrdl1*, stem cell niche factors previously shown to be expressed in the intestinal MM [10] (Fig 3B), and we annotated the *Hhip*-cells as MM-SMCs. The *Hhip+* cells, on the other hand, lacked the stem cell niche genes and instead expressed e.g., *Sncaip* and *Pdgfc* (Fig 3B). Interestingly, *Hhip* mRNA expression was limited to alpha smooth muscle actin (encoded by *Acta2*)-expressing tree-like structures within the mucosa of the colon and small intestine but was absent from the MM and MP (Fig 3C), confirming the *Hhip*-expressing cells being specific to mucosal stroma. Since they did not express fibroblast markers, we called them myocytes [9, 10]. In the integrated data, despite not clustering separately, the myocyte and MM-SMC signatures were observed in the opposite sides of the SMC cluster, highlighting their transcriptional differences (Fig 3D). In summary, the mouse SMCs clustered into two subpopulations – *Hhip*+ myocytes located in the mucosa and MM-SMCs expressing stem cell niche factors. Of note, cells from MP were not identified, likely due to their exclusion in the tissue dissociation.

**Fig 3.**
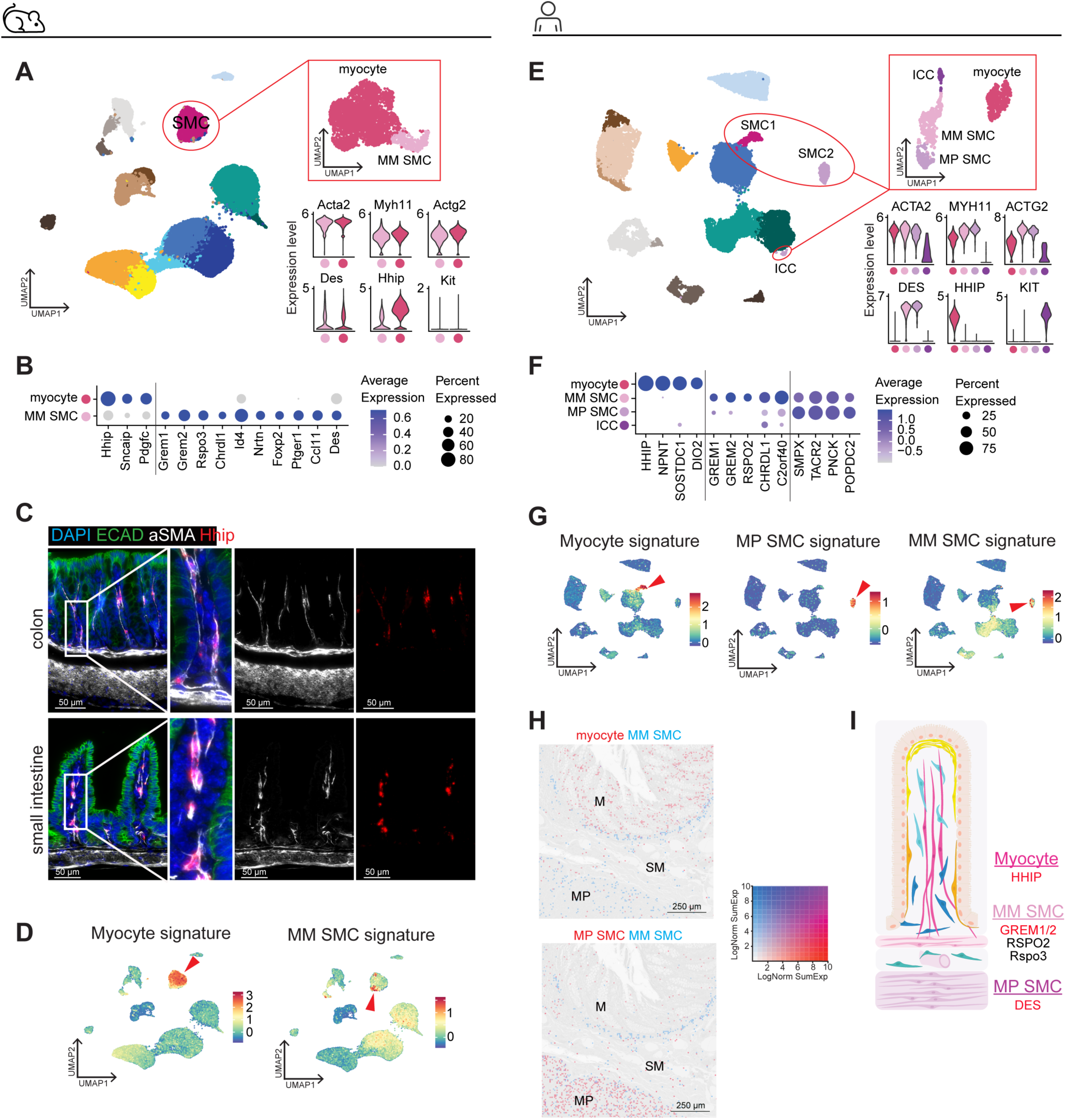
SMC subtypes in mouse and human colon. (A) UMAP of re-clustered mouse SMCs (smooth-muscle cells) and violin plots depicting indicated SMC/myofibroblast /myocyte and Interstitial cells of Cajal (ICC) markers. (B) Dot plot depicting marker genes for myocytes and MM-SMC in mice. These displayed genes are used as signatures in D. (C) mRNA *in situ* hybridization of *Hhip* and immunofluorescent staining of αSMA in mouse colon and small intestine. ECAD: E-cadherin, epithelial cells; DAPI: nuclei. (D) Feature plots of myocyte and MM-SMC signatures (from B) in mouse colon mesenchyme. (E) UMAP of re-clustered human SMCs and violin plots of indicated SMC/myofibroblast/myocyte and ICC markers. (F) Dot plot depicting marker genes for indicated cell types in human. The genes shown are used in signatures in G and H. (G) Indicated signature (from F) feature plots in integrated human colon mesenchyme data. (H) Signatures (from F) of human myocytes, MM-SMCs and MP-SMCs projected onto spatial transcriptomics data of normal human colon. M, mucosa; SM, submucosa: MP, muscularis propria. (I) Updated model of mouse and human myocytes and SMCs.

Next, we investigated the human SMC cluster by subsetting and re-clustering (Fig 3E). Consistent with the mouse analysis, the dataset included *HHIP*+ cells, interpreted as mucosal myocytes, and *GREM1+ GREM2+ RSPO2*+ cells representing MM-SMCs (Fig 3E and 3F). In addition, a third population expressing the SMCs markers, but lacking *HHIP* and the stem cell niche genes, was noted, suggesting MP origin (Fig 3E and 3F). ICCs also clustered among the SMCs but were distinguishable based on expression of *KIT* and lack of SMC markers (Fig 3E and 3F). In the whole integrated data, MM-SMCs and MP-SMCs clustered together, separately from the myocytes (Fig 3G), indicating close transcriptional similarity and distinction from myocytes. For example, *DES*, encoding for Desmin, was highly expressed in MM-SMCs and MP-SMCs but not in myocytes, indicating *DES* as a marker for non-mucosal SMCs in humans (Fig 3E).

To validate the suggested location of myocytes and SMCs in the human colon, we projected the signatures of each cell type onto spatial transcriptomics data of normal human colon [47]. As expected, myocyte signatures localized within the mucosa, MM SMCs in the muscularis mucosae, and MP SMCs in the muscularis propria (Fig 3H). Thus, our analysis provides a detailed annotation of colonic SMCs: mucosal myocytes, MM-SMCs, and MP-SMCs, together with their transcriptional and spatial characteristics (Fig 3I).

### Characterization of MIFs

The *Pdgfra*^high^ *Kcnn3*+ MIFs were identified in both mouse and human integration analyses (Fig 1B, 1D and 1F; S2H Fig). Interestingly, MIFs did not resemble any of the previously identified colon fibroblast subclusters and, despite clustering with fibroblasts and expressing *PDGFRA,* also expressed a set of unique genes absent from all other fibroblast populations (Fig 4A and 4E). Interestingly, gene set enrichment analysis (GSEA) indicated neuron-related biological functions, including voltage-gated channel activity and ion transmembrane transporter activity for MIFs (Fig 4B and 4F). Tissue staining confirmed specific KCNN3 expression in the PDGFRA+ cells exclusively in the MP in the mouse (Fig 4C) and human colon (Fig 4H), as well as in the mouse small intestine (Fig 4D), consistent with the spatial transcriptomics data (Fig 1G).

**Fig 4.**
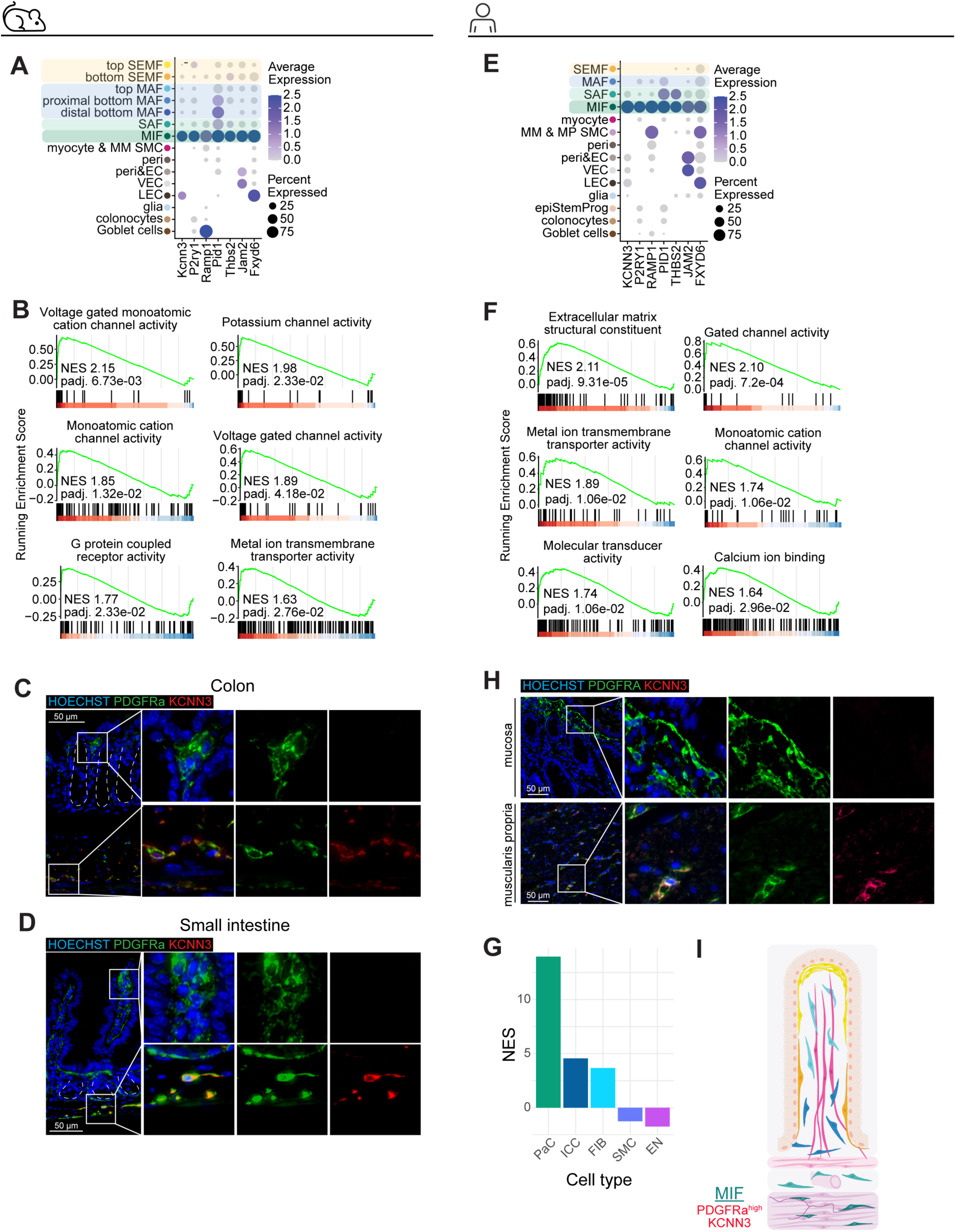
Characterization of MIFs. (A) Dot plot depicting marker genes for mouse MIFs (muscle-embedded interstitial fibroblasts). (B) GSEA of the mouse MIF signature against indicated Gene Ontology Molecular Function (GOMF) gene sets. (C) Immunofluorescent staining of PDGFRA and KCNN3 in the muscularis propria in mouse colon and (D) small intestine. Dashed line indicates crypts. (E) Dot plot depicting marker genes for indicated human cell types. (F) GSEA of human MIF signature against indicated GOMF gene sets. (G) Normalized enrichment scores (NES) of GSEA of human MIF signature against gene sets from human colon-derived cells: PDGFRA+ cells of the smooth muscle (PaC), intestinal cells of Cajal (ICC), PDGFRA+ cells of the colon mucosa (FIB), smooth muscle cells of the colon muscle (SMC), enteric neurons from colon myenteric ganglia (EN). (H) Immunofluorescent staining of PDGFRA and KCNN3 in the muscularis propria in human colon. (I) Graphical representation of MIFs in the mouse and human colon.

The neuron-related gene expression in MIFs and their clustering in the vicinity of ICCs (S2E Fig) prompted us to compare their transcriptional similarities with previously published datasets of human colon-derived cells, including 1) *PDGFRa*+ cells isolated from the smooth muscle layers, 2) ICCs, 3) *PDGFRa*+ fibroblasts isolated from the mucosa, 4) SMCs, and 5) enteric neurons [58, 59]. Notably, the MIF signature was highly enriched with the smooth muscle-derived *PDGFRa*+ cell gene set (NES >13) (Fig 4G). Thus, our analysis indicates that MIFs correspond to the previously described smooth muscle embedded PDGFRA+ fibroblasts (Fig 4I) [60–64] that have been largely missed in previous scRNA-seq analyses, presumably due to their small yield especially using mouse colon tissue dissection protocols.

### Identification and characterization of CRC CAF subsets

Having annotated the normal fibroblast heterogeneity in human colon, we next set out to characterize the cancer-associated fibroblasts (CAF) in CRC. We integrated scRNA-seq data from 62 CRC patients (both cancer and normal adjacent tissue samples) across five cohorts (CRC16, JSC, KUL3, KUL5, SMC; S1E Fig) [26, 44], resulting in a total of 327.859 cells including epithelial cells, immune cells, and mesenchymal cells (Fig 5A; S5A, S5B and S5C Fig). To focus on the mesenchymal compartment, we re-clustered the mesenchymal cells (n = 56.484), including fibroblasts, SMCs, pericytes, endothelial cells, and glial cells (Fig 5B, S5D-F Fig). *KIT*+ ICCs and *CCL19*+ reticular fibroblasts were also present in the data but did not for separate clusters (S5G Fig). Interestingly, comparison of the mesenchymal cells derived from CRC and normal adjacent colon revealed major changes within fibroblasts, where four cancer-specific clusters emerged (Fig 5C). The endothelial tip cells were also primarily derived from cancer samples, reflecting their role in vessel sprouting during tumor vascularization. However, only minor changes were noted in other mesenchymal clusters, indicating that cancer context induces major transcriptional changes specifically in fibroblasts. The cancer-specific fibroblast clusters were named CAFs, consistent with their expression of the CAF marker *FAP* [65] (Fig 5D). Notably, however, the CRC samples also included all normal fibroblast clusters, suggesting the presence of transcriptionally normal colon fibroblasts or representation of normal tissue in the resected CRC samples (Fig 5C and 5D, S5F Fig).

**Fig 5.**
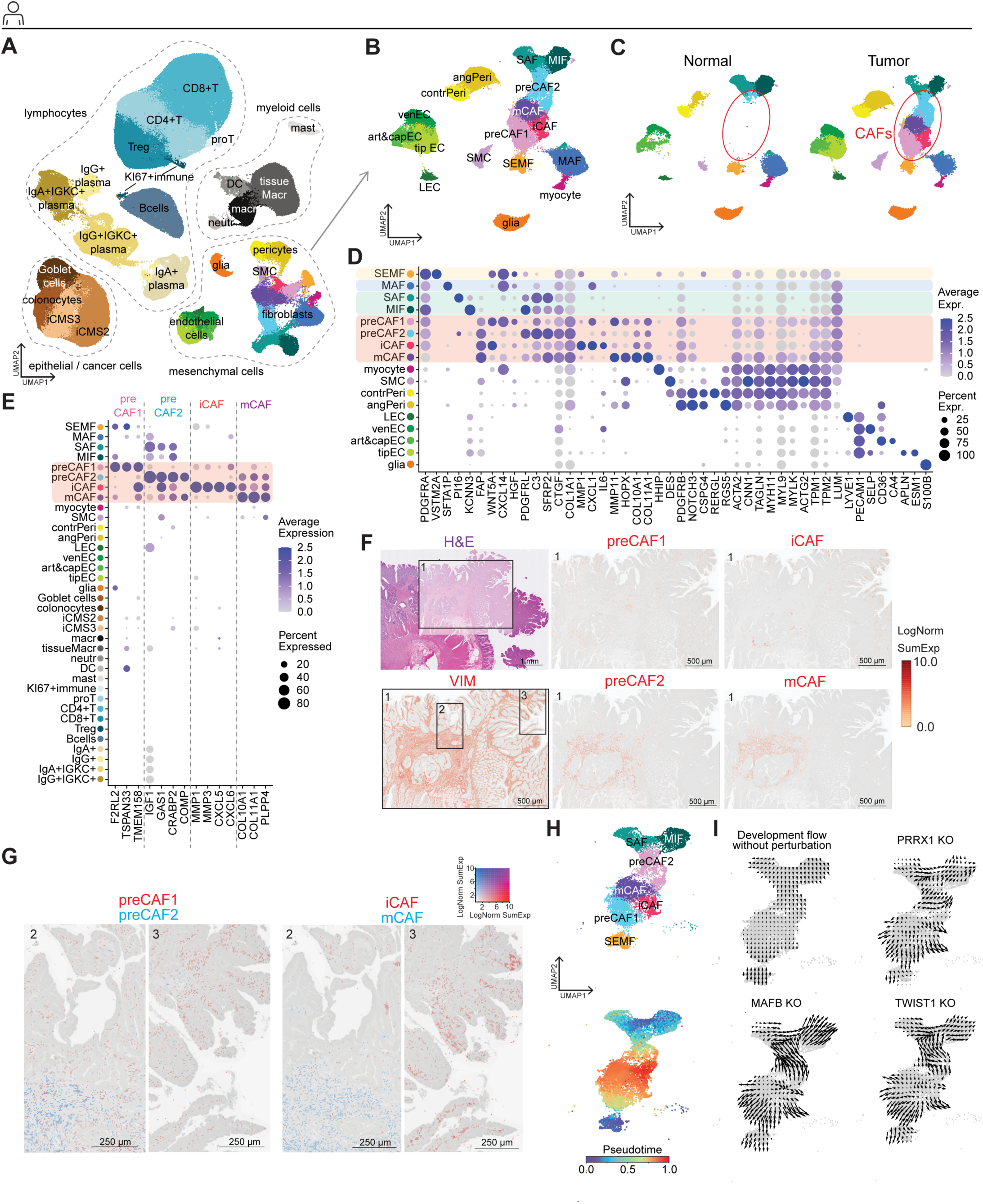
Identification of fibroblast subsets, their spatial distribution and transcriptional regulators in CRC. (A) UMAP of integrated scRNA-seq data from CRCs and normal adjacent tissue from 62 patients (n = 327.859 cells). (B) UMAP of re-clustered mesenchymal cells from A (n = 56.484). (C) UMAP shown in B split for CRC and normal adjacent tissue derived colon mesenchymal cells. (D) Dot plot depicting markers for the clusters shown in B and C. (E) Dot plot depicting CAF subtype signature genes across the whole dataset (clusters shown in A). (F) Spatial transcriptomics showing the stromal marker *VIMENTIN* (VIM) and CAF subtype signature (from E) expression in CRC tissue. (G) Zoom-ins of areas 2 and 3 (from F) with overlayed preCAF1 and preCAF2 signatures (left panels) and iCAF and mCAF signatures (right panels). (H) Fibroblast subtypes used for computational transcription factor knockout simulations. The pseudotime was calculated by allowing cells to take two simultaneous differentiation trajectories (from MIF to mCAF; from SEMF to iCAF). (I) Vector fields describing the development of the cellular fibroblast system without perturbation (control, gradient of pseudotime), or after simulated transcription factor knockouts. contrPeri, contractile pericytes; angPeri, angiogenic pericytes; venEC, venular endothelial cells; art&capEC, arteriolar and capillary endothelial cells; tipEC, endothelial tip cells, iCMS intrinsic consensus molecular subtype of CRC; macr, macrophages; tissueMacr, tissue-resident macrophages; neutr, neutrophils; DC, dendritic cells; mast, mast cells; KI67+ immune, proliferative immune cells; proT, proliferative T cells; Treg, regulatory T cells; Bcells, B cells; IgA+, IgA+ plasma cells; IgG+, IgG+ plasma cells; IgA+ IGKC+, IgA+ IGKC+ plasma cells; IgG+ IGKC+, IgG+ IGKC+ plasma cells.

Next, we investigated the CRC CAF subsets in more detail. One of the CAF populations specifically expressed *MMP1, CXCL1,* and *IL6* as well as other inflammatory genes, closely aligning with the previously described inflammatory CAFs (iCAFs) [66–69] (Fig 5D, S6A Fig). We also identified a cluster expressing the myofibroblast/matrix CAFs (mCAFs) genes, such as *MMP11, HOPX, COL10A1,* and *COL11A1* (Fig 5D, S6B Fig) [67, 70]. The two remaining CAF populations lacked clear specific markers and shared transcriptional features with both normal-like fibroblasts and iCAFs/mCAFs, suggesting transitional states, and were named precursor (pre) CAFs. Namely, the preCAF1 cluster shared gene expression with SEMFs and iCAFs, (Fig 5D, S6C Fig), while the preCAF2 cells shared features with SAFs/MIFs and mCAFs (Fig 5D, S6D Fig). These observations prompted a hypothesis that preCAF1 and 2 may represent transitional states towards iCAFs or mCAFs, respectively, with different cells of origin: SEMFs for preCAF1 and SAFs/MIFs for preCAF2.

Next, we addressed the correlation of the CAF subset signatures (Fig 5E) to CRC patient survival probability. Interestingly, preCAF2 and mCAF signatures were correlated with reduced probability of overall survival, while a trend towards improved survival was observed for preCAF1 and iCAF signatures (S6E Fig). To explore the spatial distribution of the identified CAF subtypes, we projected their signatures (Fig 5E) onto CRC spatial transcriptomics data [47]. The preCAF1 and iCAF gene expression was found in the vicinity of cancer cells around the tumor edges, consistent with their suggested origin from SEMFs (Fig 5F and 5G). In contrast, preCAF2s and mCAFs were mainly located deeper in the tumor stroma, within the invasive front, consistent with SAF/MIF origin (Fig 5F and 5G). Importantly, the CAF signature genes were barely detectable in normal colon tissue, confirming their cancer-specificity (S7A Fig). Normal fibroblast signatures were, however, also observed in CRC tissue, suggesting that not all fibroblasts in CRC represent altered fibroblast states, consistently with our scRNA-seq analysis (S7B and S7C Fig).

In accordance with the transcriptional and spatial differences, trajectory and pseudo-time analyses using the Monocle3 algorithm [71] suggested that iCAFs and mCAFs develop via separate trajectories from distinct tissue-resident fibroblast subsets: SEMFs being the cells of origin for iCAFs through a preCAF1 state, and SAFs/MIFs polarizing into mCAF phenotype via preCAF2s (S7D-G Fig). To further explore gene regulatory changes during the CAF differentiation, we constructed gene regulatory networks (GRNs) for each identified fibroblast subtype and performed a gene ranking in which genes positioned across control points within the GRNs received a high score based on betweenness centrality (S7D Fig). Among the top-ranked genes, 22 were transcription factors (TFs). Next, we employed computational knockout simulations to assess how perturbations of these TFs would affect CAF differentiation. These simulations included the four CAF populations and normal fibroblast subtypes predicted to give rise to CAFs based on the Monocle3 analysis (SEMFs, SAFs, and MIFs) (S7E-H Fig), allowing them to undergo simultaneous differentiation along both identified trajectories towards iCAF and mCAF (Fig 5H). Using this approach, we identified *PRRX1*, *MAFB,* and *TWIST1* as potential TFs regulating CRC CAF differentiation (Fig 5I). Consistently, the expression levels of the candidate TFs were highest in CAFs (S7H Fig).

Next, we investigated whether the relative abundances of the CAF subsets correlated with molecular and clinical features across CRCs (n = 40, S8A and S8B Fig). On average, 34.7 % of CAFs across these samples represented preCAF1; 14.1 % were iCAFs; 24.4 % preCAF2; and 26,8 % mCAFs (S9A Fig). Despite the relatively low numbers of samples falling to each category (n = 6 - 11 per group), we correlated the CAF subtype abundance across the CRC CMS 1-4 classification [23] (S9B and S9C Fig). Interestingly, the CMS2, the canonical subtype, presented with a significantly higher abundance of preCAF2s as compared to other subtypes (S9D Fig). Similarly, the CMS3, the metabolic subtype, showed a significantly (p < 0.05) higher abundance of preCAF1s (S9E Fig). Additionally, a tendency towards increased proportion of mCAFs in CMS4 was observed (S9F Fig). No consistent differences in CAF proportions between the iCMS2 and iCMS3 subtypes [44] or right-left sidedness were noted (S9G-J Fig), but rectal tumors presented with a higher abundance of preCAF1s as compared to colonic tumors (S9K and S9L Fig). MSI-H status showed a trend towards increased proportion of preCAF2, while iCAFs were significantly more abundant in early (1-2) versus late stage (3-4) CRC (S9M-R Fig).

We then examined the correlation of CAF proportions with the mutational status for *APC*, *KRAS*, *TP53, BRAF,* and *PIK3CA* (S10 Fig). Significant (p < 0.05) or borderline significant (p = 0.055 - 0.066) correlations were found with increased abundance of mCAFs in KRAS mutant cancers, and preCAF1s in *TP53* mutant and *PIK3CA* wild-type cancers (S9B and S9C Fig, S9f and S9G Fig, S9I and S9J Fig). Moreover, mCAFs were significantly enriched in late-stage *KRAS*-mutated cancers (S10D and S10E Fig).

### CTHRC1 represents a pan-CAF marker in CRC

Multiple genes have been suggested as specific pan-CAF markers in CRC and other cancers, including *ACTA2, PDGFRA, PDGFRB, FAP, S100A4, PDPN, LRRC15, POSTN* or *TNC* [72, 73]. Of these, only *FAP* faithfully marked CAF populations in our analysis (Fig 6A). In addition to *FAP*, we identified *CTHRC1*, *INHBA*, *THBS2* and *WNT2* as the most specific and sensitive markers for CRC CAFs. Projecting the pan-CAF signature (*CTHRC1*, *INHBA*, *THBS2, WNT2, FAP)* (marked by asterisk in Fig 6A) onto CRC spatial transcriptomics data [47] showed clear signal in the tumor stroma but was barely detected in normal tissue as predicted (Fig 6B, S11A Fig). Following our data suggesting *CTHRC1* as a pan-CAF marker in CRC, CTHRC1 protein staining was noted in mesenchymal cells in 96% of CRCs (n = 279/292) and was not detected in normal colon mesenchyme (n = 0/4) (Fig 6C and 6D, S11B). The extent of CTHRC1 staining did not, however, affect overall survival in this cohort (S11C Fig).

**Fig 6.**
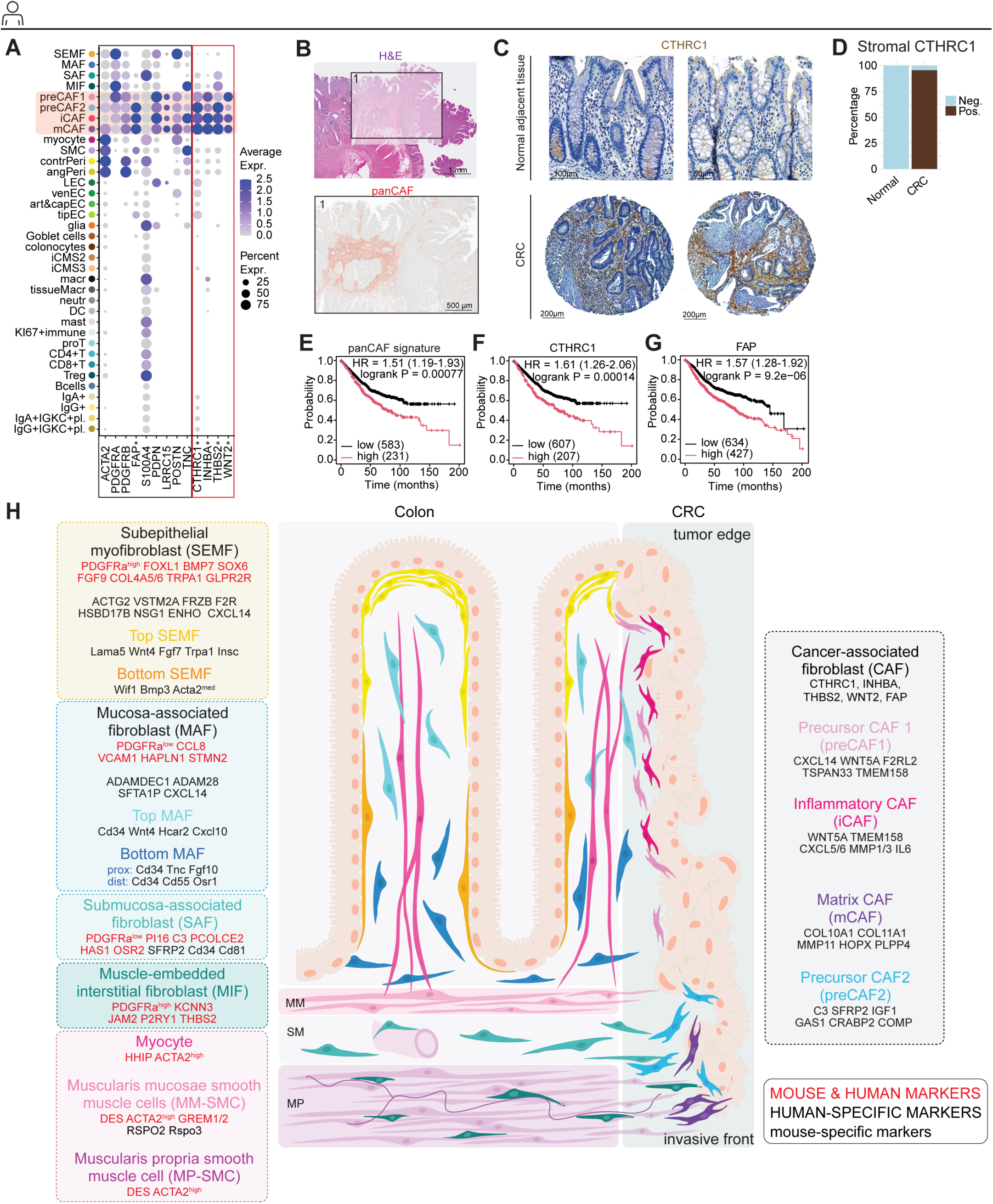
Identification of CTHRC1 as a pan-CAF marker in CRC. (A) Dot plot depicting previously proposed CAF markers (left, black box) and novel CRC pan-CAF markers identified in this study (right, red box). (B) CRC pan-CAF signature (*CTHRC1*, *INHBA*, *THBS2* and *WNT2*, from A) projected onto spatial transcriptomics data of CRC tissue. (C) Representative images of CTHRC1 staining in normal adjacent colon and CRC. (D) Quantification of CTHRC1 staining in normal adjacent colon and CRC mesenchyme. (E-G) Kaplan-Meier plots depicting the correlation between mRNA expression of the panCAF signature (*CTHRC1, INHBA, THBS2, WNT2, FAP*), as well as *CTHRC1* and *FAP* alone with overall survival probability in CRC. mRNA-sequencing data derived from: GSE12945, GSE12294, GSE14333, GSE123985, GSE17538, GSE10800, GSE26682, GSE20540, GSE21595, GSE33114, GSE34489, GSE37892, GSE38832, GSE39582, GSE41258, GSE92921. (H) Schematic representation and specific markers for the identified mouse and human colon fibroblast and SMC subpopulations (left) and CRC CAF subsets with suggested trajectories (right).

Since the abundance of CAFs appears to be increased in late-stage CRC (S11D Fig) and we found both *CTHRC1* and *FAP* highly expressed in CMS4 subtype of CRCs (S11E and S11F Fig) [74], we set out to investigate the effects of these markers on survival between early and late-stage CRC and across CMS statuses in multiple published RNAseq datasets. While the pan-CAF signature, as well as *CTHRC1* and *FAP* expression alone, correlated with poor overall survival in CRC (Fig 6E-G), the effects were specific to late-stage CRC (S11G-I Fig) and CMS4 patients (S11J-L Fig). Moreover, *CTHRC1* mRNA expression alone reflected the survival predictions obtained using the pan-CAF signature and *FAP,* indicating that *CTHRC1* mRNA expression alone is a predictor of overall survival in late-stage and CMS4 CRC. In summary, these results suggest that both the CAF signature expression and the pan-CAF marker *CTHRC1* expression alone are associated with reduced survival probability in late-stage and CMS4 CRC subtypes. Thus, CAFs indeed represent intriguing therapeutic targets.

### Refined high-resolution model for colon fibroblast heterogeneity in homeostasis and CRC

Finally, we used our detailed analysis to construct a systematic and transparent summary of the fibroblast populations in both healthy colon and CRC (Fig 6H). We also compared our results with previously published data by matching the clusters using overlapping markers. Despite the very heterogeneous nomenclature, the reported gene expression patterns allowed us to match most of the previously proposed normal colon fibroblast (S3 Table) and CRC CAF subtypes (S4 Table) with the clusters identified in our analysis. Of note, our analysis indicates that the previously suggested CRC CAF subsets represented in addition to CAFs also cell types such as normal fibroblasts, pericytes, and SMCs. For example, we did not find a CAF cluster matching the proposed apCAFs (*CD74*+) [33, 35, 70, 75, 76], which may instead represent reticular fibroblasts (S5G Fig). We also failed to identify a separate pCAF (*MKI67*+) cluster [33, 42, 68]. Finally, to support easy comparison of our data and other studies, we provide an easy-to-use comparison of our annotation with previously suggested nomenclature, along with key marker genes and spatial information (Table 1).

**Table 1.**
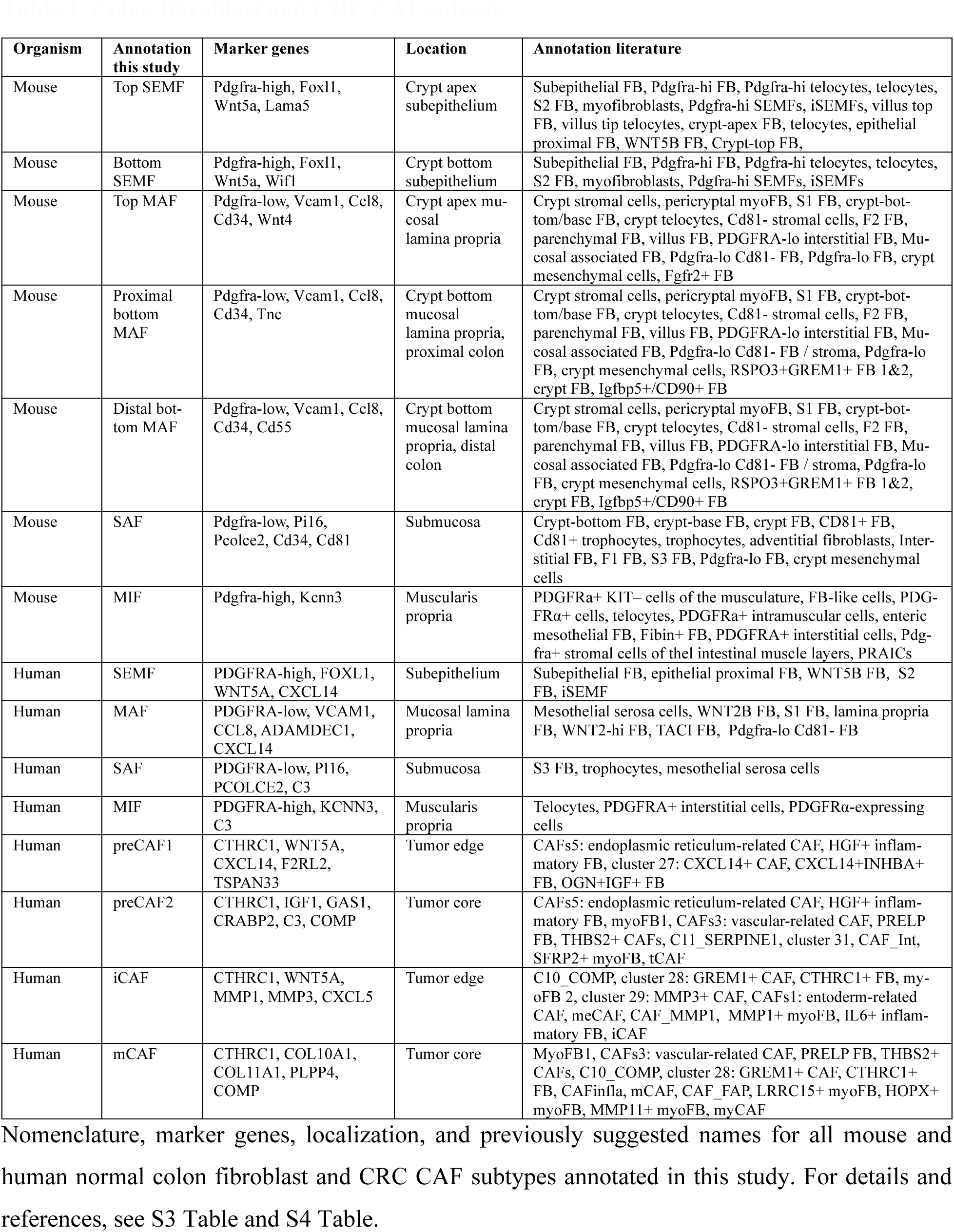
Colon fibroblast and CRC CAF subsets.

## DISCUSSION

Recent discoveries highlighting fibroblasts as major regulators of tissue maintenance and pathological processes, including cancer, have fueled the need to understand fibroblast heterogeneity in healthy and pathogenic tissues [12, 77–81]. This study aimed to (1) generate a high-resolution map and annotation of mouse colon fibroblast subclusters; (2) characterize human colon fibroblasts using benchmarking data from mice and spatial transcriptomics; and (3) identify and annotate CRC CAF subsets and investigate their potential differentiation trajectories and association with clinical parameters.

Several studies have addressed fibroblast heterogeneity in mice, resulting in a consensus view of the three major subclasses [4–11] while the nomenclature remains variable (S3 Table). The heterogeneity of human colon fibroblasts is less well established. Our side-by-side integrated analyses of mouse and human colon fibroblasts, followed by spatial validation, allowed us to establish a coherent, location-based nomenclature using previously proposed (SEMF, MAF) [8] and novel terms (SAF, MIF). Importantly, we found remarkable similarities between mouse and human colonic fibroblast organization and provide both shared and species-specific markers for the identified fibroblast subsets, allowing easy comparison with other datasets (Fig 6H, Table 1). It is likely that the fibroblast identity represents a cell state, rather than cell type, and is largely regulated by the tissue context. Indeed, PI16-expressing fibroblasts have been proposed to be associated with vasculature and in colon, PI16 expression localizes close to the endothelium, suggesting that the proximity of endothelial cells may be essential for the SAF phenotype [82]. Moreover, fibroblast subtypes have been shown to be re-established at correct locations after tissue damage [83]. Interestingly, though, cell culture experiments addressing the functional properties of isolated primary fibroblast subsets suggest at least some degree of stability in short-time cell culture models [4, 6, 9, 10, 51]. Establishing conditions that maintain the fibroblast state for extended periods *ex vivo* would benefit the functional characterization of the identified fibroblast subsets.

Our study complements the existing knowledge of mouse colon fibroblasts subsets by showing that SEMFs and MAFs further divide into distinct subpopulations, reflecting gradients along the crypt-apex (basal-luminal) and proximo-distal axis. Notably, similar observations have been reported before in the small intestine and colon [4, 11]. The crypt-apex gradient appears to take place also in the mouse small intestine, where the low representation of top SEMF and MAF subtypes and a lack of MIFs likely stems from the commonly used protocols, where both the intestinal villi and muscle layers are excluded in preparation of single cell suspensions. Our analysis suggests that human fibroblasts do not form similar subpopulations, but it remains possible that our dataset lacked sufficient power to reveal further subclusters.

SMCs are characterized by the expression of genes such as *ACTA2*, *ACTG2*, *MYH11,* and *DES*, while the cells co-expressing SMC markers and *HHIP* are often referred to as myofibroblasts [2, 4, 5, 13, 43, 56, 57, 84]. In our analysis of the normal colon, SEMFs were the only cell type with a clear fibroblast identity (clustering with other fibroblasts, expressing *PDGFRA*), together with expression of the SMC markers *Acta2* (in mice) and *ACTG2* (in humans), and could therefore be classified as myofibroblasts. Since the *Acta2*^high^ *Hhip*+ cells lacked fibroblast markers but were transcriptionally and spatially distinct from MM-SMCs and MP-SMCs, we propose the term myocyte for these cells, consistent with suggestions in the small intestine and stomach [9, 10]. The myocytes are located in the mucosal core of the colonic and intestinal lamina propria, and *HHIP* encodes for a Hedgehog signaling inhibiting factor. Since the intestinal and colonic epithelium express Hedgehog ligands [85], we hypothesize that the surface-core gradient of Hedgehog signaling could contribute to mucosal SEMF and MAF identity. Interestingly, myocytes also express *Pdgfc,* a PDGFRA ligand and a factor regulating endothelial growth. Thus, myocyte-derived PDGFC could play a role in programming the MAF and SEMF identities either directly or indirectly by regulating vascular growth, which affects oxygen abundance. Indeed, it was recently reported that hypoxia induces expression of WNT5A (SEMF marker) and an inflammatory fibroblast phenotype in the edges of colonic tumors [86], indicating that the SEMF identity might be driven, at least partly, by hypoxia.

Interestingly, we observed a *PDGFRA*^high^ *KCNN3*+ population clustering with fibroblasts and localizing within the muscularis propria and therefore named them MIFs. We propose that MIFs represent the fourth major colon fibroblast subset but were missed in previous mouse scRNA-seq experiments due to the removal of the muscle layers in the cell dissociation protocols. The corresponding human cluster was more prominent, and the human datasets also contained cells from muscularis propria, reflecting the inclusion of the muscle wall in CRC resections. Thus, the sampling method (e.g., biopsy vs. resection) is an important aspect to consider when comparing fibroblast compositions across different samples. MIFs likely correspond to the recently reported PDGFRA^high^ interstitial cells (PRAICs) of the mouse small intestine, although the marker genes for PRAICs were not reported [10]. Interestingly, other methods have revealed PDGFRA and KCNN3-expressing cells in the vicinity of ICCs throughout the gastrointestinal tract smooth muscle layers [60–64]. Those cells have been suggested to participate in the regulation of SMCs in concert with ICCs and neurons [87–91]. Thus, we link the scRNA-seq findings with a previously characterized cell type and suggest the *PDGFRA*^high^ *KCNN3*+ cells represent fibroblasts that display neuronal properties and likely act as regulators of the smooth muscle layer functions, such as peristalsis.

In CRC, CAFs have emerged as key players and drug targets, mandating a detailed understanding of their heterogeneity. Tens of different names have been proposed for suggested CAF subsets (S.Table 4), but a coherent understanding has not been established. This is in part due to the incomplete understanding of the normal human colon fibroblasts and other mesenchymal cell subtypes. Our results indicate that the scRNA-seq analyses performed to date identify rather reproducible cell types/states, but only a subset of the proposed populations represents CAFs (S.Table 4). Importantly, our analysis revealed only four cancer-specific (CAF) clusters but showed that all the normal fibroblast subtypes persist within CRC. Interestingly, we identified iCAFs and mCAFs – the most well-characterized CAF subtypes – which expressed similar marker genes observed in corresponding CAF populations in other tissues [66–70]. However, in contrast to pancreatic cancer, where iCAFs were shown to localize at a distance from cancer cells and be associated with poor survival [92], in CRC, iCAFs are seem to be situated closer to the cancer cells, while mCAFs reside closer to the invasive front, and the mCAF signature expression predicts reduced survival probability. This is consistent with the recent report indicating expression of WNT5A in the tumor edge in iCAF-like cells that derive from BMP-expressing normal fibroblasts [86]. The other two CAF populations, preCAF1 and preCAF2, shared markers with SEMFs and SAFs/MIFs, respectively, and were interpreted as transitional states. Indeed, our trajectory and pseudotime analyses indicated that iCAFs develop from SEMFs via preCAF1s, and mCAFs arise from SAFs/MIFs through a preCAF2 state. Thus, only two functionally different CAF subtypes/trajectories may exist in CRC. Additionally, our *in silico* knock-out simulations identified three transcription factors, *PRRX1, MAFB,* and *TWIST1*, which showed potential regulatory functions in CAF differentiation. While the roles of MAFB and TWIST1 in CRC CAF differentiation are not yet characterized, PRRX1 has previously been shown to regulate CRC CAFs, supporting the validity of our data-driven approach for identifying transcriptional regulators critical for fibroblast regulation during colon cancer [93]. Thus, these genes represent an intriguing avenue of research for the future.

The cellular origin of CAFs in CRC remains under debate, with different studies proposing tissue-resident fibroblasts and other cells such as bone marrow-derived mesenchymal stem cells [94], SMCs [95], pericytes [96], endothelial cells [97], epithelial cells [98], and adipocytes [99, 100] as sources of CAFs. Indeed, pericytes are commonly annotated as fibroblasts or CAFs in the CRC field despite their distinct transcriptional signature (*RGS5, NOTCH3, PDGFRB, NDUFA4L2*) [101–104]. In our analysis, pericytes were transcriptionally distinct from the CAF populations. Moreover, CAFs clustered with fibroblasts, and the trajectory analysis suggested a fibroblast origin. Thus, we propose that the majority of CAFs originate from tissue-resident fibroblasts, and that some of the previously identified subsets such as vascular CAFs [39, 68, 75, 105] may represent pericytes and CRC apCAFs [33, 35] represent reticular fibroblasts that also express high levels of *CD74* (S5G Fig). Additionally, we did not identify a distinct *MKI67*+ proliferative pCAF [33, 42] population (S5G) and hypothesize that while apCAFs and pCAFs may be represented in some cancer types, they do not form a specific entity in CRC. Of note, contributions from circulating mesenchymal stem cells cannot be excluded by our analysis.

The different CAF subset proportions between CRC CMS subtypes and genetic features suggest that fibroblast signatures may underlie some of the CMS classifications, and that different mutations could influence the CAF subtype composition, placing fibroblast phenotypes as important determinants of patient survival and treatment efficacy. Interestingly, our results suggest an increased abundance of mCAFs in CMS4 patients, commonly known to display a fibroblast-rich tumor phenotype and poor patient survival. Additionally, *KRAS* mutations were associated with high mCAF proportion in late-stage CRC. Consistently, a previous study in pancreatic cancer suggested that targeting mutant *KRAS* in cancer cells reprograms CAFs towards normal-like fibroblasts, raising the possibility that targeted therapies might also alter CAF subtype composition in CRC [106]. Thus, we hypothesize that targeting CAFs, and especially mCAFs, could benefit CRC patients belonging to the CMS4 subgroup and/or carrying the epithelial *KRAS* mutation, especially in late-stage CRC. It will be interesting to address in larger datasets if CAF composition may drive transcriptional differences between the CMS subtypes, and whether specific composition of mutations could play a role in the CAF reprogramming in CRC.

Specific and sensitive markers that differentiate CRC CAFs from normal fibroblasts could serve as valuable biomarkers or drug targets. Our analysis revealed that among previously suggested CAF markers, only *FAP* was expressed across CAFs and absent from other cell types. Instead, our analysis identified *CTHRC1*, *INHBA*, *THBS2,* and *WNT2* as CAF-specific markers. Interestingly, tissue staining confirmed that CTHRC1 protein expression was specific to the cancer stroma. CTHRC1 plays a crucial role in ECM remodeling by limiting collagen deposition and enhancing ECM plasticity, which are reported cancer-promoting functions driven by CAFs [107–111]. Therefore, CTHRC1 might be functionally linked to tumor progression by facilitating cancer cell migration and angiogenesis. Given its secreted nature and expression already in the preCAFs, CTHRC1 could potentially serve as a biomarker for early-stage CRC and thus represents a compelling target for further investigation. Interestingly, *CTHRC1* was previously suggested to mark a specific CAF subset in CRC [32, 112]. Our analysis suggests that *CTHRC1+* fibroblasts represent all CAFs, while the other clusters represent normal tissue-resident fibroblasts.

In summary, our integrative analysis enabled the construction of a high-resolution model for fibroblast heterogeneity with consistent location-based nomenclature for both mouse and human colon fibroblasts, incorporating both previously validated and newly discovered marker genes. Additionally, our study provides a systematic characterization of CRC CAF subsets, offering a standardized framework for future studies into the colon fibroblast heterogeneity in both health and disease.

## MATERIALS AND METHODS

### Mice

Two C57Bl/6 mice (one male and one female) were used for the isolation of stromal cells for single-cell RNA sequencing (scRNA-seq) from the colon and small intestine. Mice were bred and maintained in the animal house facilities of BSRC Alexander Fleming under specific pathogen-free conditions, and experiments were performed in accordance with current European and national legislation (approved by BSRC Fleming’s Institutional Committee of Protocol Evaluation in conjunction with the Veterinary Service Management of the Hellenic Republic Prefecture of Attika). Male and female C57BL/6JRccHsd mice (Mus musculus) were used for tissue collection for the validation experiments. These animals were housed at the University of Helsinki Laboratory Animal Center according to national and international legislation and guidelines (license: KEK23-006, approved by the Research Ethics Committee on Animal Research). Mice had free access to food and water.

### Isolation of intestinal stromal cells by fluorescence-activated cell sorting (FACS)

Isolation of intestinal stromal cells was performed as previously described [83]. Briefly, intestinal tissues were dissected, opened longitudinally, and washed with HBSS (Gibco) supplemented with antibiotic-antimycotic solution (Gibco). Epithelial cells were removed by incubating the tissue with 5 mM EDTA and 1 mM DTT in HBSS for 25 minutes at 37°C. The remaining tissues was pooled and digested in 300 U/mL Collagenase XI (Sigma-Aldrich) and 1 mg/mL Dispase II (Roche) in DMEM for 40-60 min at 37°C. The cell suspension was centrifuged, washed three times in FACS buffer (5% FBS (Biowest) in PBS), and the cells were counted. 1-2 million cells/100 μL were incubated with the APC-Cy7 conjugated anti-CD45 (clone 30-F11, BioLegend) and the APC-efluor780 conjugated anti-CD326 (clone G8.8, eBioscience) antibodies. Propidium Iodide (Sigma) was used for live-dead cell discrimination. Cell sorting was performed using the FACSAria III cell sorter (BD) and the FACSDiva (BD) software.

### Single-cell RNA library preparation and sequencing

100.000 FACS-sorted stromal cells from the colon and the small intestine were used fresh for the preparation of two single-cell RNA-sequencing libraries according to the 10X Chromium Single Cell 3’ v3.1 protocol and sequenced using the DNBSEQ-G400 sequencer (PE100) (BGI Genomics). FASTQ files containing raw reads were initially processed, including read alignment, unique molecular identifier (UMI) counting, filtering of empty droplets, and the generation of sample-specific count matrices, using the 10X Genomics CellRanger pipeline (version cellranger-7.1.0) function ‘cellranger count’, at default settings. Read pairs were aligned to the mouse reference genome (mm10, 10X-reference genomes, v2020). The number of read pairs assigned to each library and the estimated cell numbers were 237,166,873 bp and 6,990 cells for the colon and 210,226,057 bp and 11,576 cells for the small intestine, respectively. Original raw and processed scRNAseq data has been deposited at the Gene Expression Omnibus under the accession number GSE296873.

### ScRNA-seq data analysis

All scRNA-seq data in this study was analyzed using the Seurat algorithm V4 [113] in R (version 4.2.3). All Seurat objects were created by setting the minimum number of cells per sample to 3 and the minimum number of genes expressed per cell to 200. All objects were normalized using SCTransform.

Mouse colonic scRNA-seq data produced for this publication was filtered using nFeature RNA 1000<n<4000, nCountRNA 1000<n<13.000, percent.mt<6, percent.rb 3<n<25 and the small intestinal data was filtered using nFeature RNA 500<n<5000, nCountRNA 750<n<15.000, percent.mt<5.5, percent.rb 2.5<n<20. Variable regression included %mt and %rb. The data was clustered (graph-based Louvain clustering) using 20 and 25 principal components (PCs) and resolution 0.5 and 0.4 for colon and small intestine, respectively. For further analysis, *Pdgfra*+ fibroblasts and *Myh11*+ SMCs were subsetted and re-clustered using 15 and 25 PCs and a resolution of 0.15 and 0.3 for colon and small intestine, respectively.

Published scRNA-seq data from healthy mouse colon was obtained from the GEO database, quality controlled (QC), and integrated with the scRNA-seq data produced for this study (S1 Table) [4, 5, 7, 8]. The data was filtered according to S1 Table. Variable regression included %mt, %rb, and cell cycle scores. The data was integrated using the Seurat anchor-based integration workflow: top 3.000 variable features, RPCA, and k.anchor 7. The integrated data was clustered (graph-based Louvain clustering) using 30 principal components (PCs) and resolution 0.7. Based on low overall gene expression (nFeatureRNA), low RNA counts (nCountRNA), high ribosomal RNA expression (%rb), and high expression of *Ptprc* (CD45), two small populations were filtered out, and the data were re-clustered using 15 PCs and resolution 0.6. Cell type annotation was done using validated marker genes (Fig 1B, S2B Fig).

Human scRNA-seq data were obtained from synapse.org under the project ID syn26844071 [44]. For analyzing homeostatic human colon tissue, only samples from normal adjacent tissue were included, and lymph node data was filtered out. The data had already been quality controlled by the original authors. Further QC and filtering were done on epithelial data using a cutoff for nFeatureRNA at 6.000 and nCountRNA at 25.00,0 and on stromal data using a cutoff for nFeatureRNA at 4.500 and nCountRNA at 35.000. The data was integrated using the Seurat anchor-based integration workflow: top 3.000 variable features, RPCA, and k.anchor 10. The integrated data was clustered (graph-based Louvain clustering) using 25 PCs and resolution 0.5. One sample (patient CRC2786) displayed features of tumor samples and was thus filtered out, resulting in a total of 36 patients. For further analysis, epithelial and immune cells were filtered out, and the data was re-clustered using 20 PCs and resolution 0.4. Cell type annotation was done using validated marker genes (Fig 1D, S2D Fiig).

For the integrative analysis of human normal adjacent colon tissue and CRC tumor tissue, data were obtained from synapse.org under the project ID syn26844071 [44]. Lymph node data was filtered out. The data had already been quality controlled by the original authors. Further QC and filtering were done on epithelial data using a cutoff for nCountRNA at 50.000 and on stromal data using a cutoff for nFeatureRNA at 5.500 and nCountRNA at 70.000. The sample from patient CRC278 was filtered out to match the normal tissue analysis. This resulted in samples from 62 patients. The data was integrated using the Seurat anchor-based integration workflow: top 3.000 variable features, RPCA, and k.anchor 10. The integrated data was clustered (graph-based Louvain clustering) using 30 PCs and resolution 0.5. Cell type annotation was done using validated marker genes (Fig 5D, S5B Fig). For the mesenchymal cell analysis (Fig 5B-D), only samples with a minimum of 50 cells in a minimum of 5 different stromal cell clusters were included and the small PECAM1 and RGS5 double-positive cluster likely representing doublets formed of endothelial cells and pericytes was filtered out. This resulted in 53 patients (S5F and S5G Fig). Mesenchymal cell type annotation was done using validated marker genes (Fig 5D). For CAF abundance analyses, only the four CAF populations were subsetted from the mesenchymal analysis, and samples with a minimum of 50 CAFs in total were included. This resulted in 40 patients and a total of 15.022 cells (S8A and S8B Fig). CAF abundances were compared using the Wilcoxon rank-sum test. Variability is represented as 1.5 times the interquartile range (IQR), as shown by the whiskers in box plots.

### Spatial transcriptomics data analysis

Spatial transcriptomics data (Visium HD, 10x Genomics) from normal human colon and CRC tumors were obtained from https://www.10xgenomics.com/products/visium-hd-spatial-gene-expression/dataset-human-crc [47]. Analysis was executed in the Loupe Browser 8 app. Marker gene signatures (lists of genes, S.Table 2) were projected onto the spatial data and visualized using the following settings: Combination: FeatureSum; Scale value: LogNorm; Saturation: −100; Opacity: 30%, and images exported as tiffs.

### Immunofluorescent staining of mouse tissue

Small intestinal and colon tissues were collected, washed with PBS, and fixed in 4% paraformaldehyde at 4°C for 24 h. The tissues were embedded in paraffin as Swiss rolls with a Sakura Tissue-Tek VIP 5 Jr tissue processing system. Then, 5 µm sections were cut using a microtome (Shandon AS325, Marshall Scientific). Paraffin-embedded sections were deparaffinized in a xylene and ethanol series. Antigen retrieval was performed using 1× Dako antigen retrieval solution at pH 9 (S2367, Agilent Technologies) at 95°C for 25 min, followed by a 30 min incubation at room temperature (RT). Samples were washed 2 x 5 min with PBS, and permeabilization was performed with 0.1% Triton X-100 in TBS (TBS-T) for 10 min at RT, followed by blocking for 30 min using TNB blocking buffer. The samples were incubated with anti-PDGFRa (AF1062, R&D systems, 1:100) and anti-KCNN3/KCa2.3 (APC-025, Alomone Labs, 1:400) antibodies overnight at 4°C in blocking buffer. The next day, samples were washed 5 min with TBS containing 0.1% Tween-20 and 2 x 10 min with PBS. Then, samples were incubated with secondary antibodies [AlexaFluor 546 donkey anti-goat IgG (A11056, Thermo Fisher Scientific) and AlexaFluor 647 donkey anti-rabbit IgG (A31573, Thermo Fisher Scientific)] were incubated in blocking buffer (1:500 dilution) for one hour at RT and washed 3 x 5 min in TBS containing 0.1% Tween-20. Nuclei were stained with 10 µg/ml Hoechst 33324 (62249, Thermo Fisher Scientific) in PBS for 3 min. Slides were rinsed with TBS and water before mounting with Immu-Mount (9990402, Thermo Fisher Scientific).

### Immunofluorescent staining of vibratome sections from mouse tissue

The mice were perfused with 15 ml of PBS, followed by 15 ml 4% paraformaldehyde. Small intestinal and colon tissues were collected as a whole, washed with PBS, and flushed and fixed with 4% paraformaldehyde overnight at 4°C. Tissues were cut into 1 cm pieces and embedded in 3 % TopVision Low Melting Point Agarose in PBS. Then, 70-100 µm sections were cut using a vibratome (Vibratome Series 1000 – Tissue Sectioning System) and stored in 5 % Na-Azide in PBS. Permeabilization was performed with 0.1% Triton X-100 in PBS (PBS-T) for 3 x 5 min at RT, followed by blocking for 1 h using 0.3% PBS-T containing 3 % goat serum and 3 % donkey serum. The samples were incubated with anti-PDGFRa (#3174, Cell Signaling, 1:100) and anti-TNC (MAB2138, R&D Systems, 1:200) antibodies overnight at 4°C in 0.3% PBS-T containing 1 % goat serum and 1 % donkey serum. Samples were then washed 3 x 15 min with 0.1 % PBS-T and incubated with secondary antibodies [AlexaFluor 594 goat anti-rat IgG (A11007 Thermo Fisher Scientific, 1:500) and AlexaFluor 647 donkey anti-rabbit IgG (A31573, Thermo Fisher Scientific, 1:500)] in 0.3% PBS-T containing 1 % goat serum and 1 % donkey serum for 1 h at RT and washed 3 x 10 min in 0.1% PBS-T. Nuclei were stained with 10 µg/ml Hoechst 33324 (62249, Thermo Fisher Scientific) in PBS for 10 min at RT. Samples were washed 3 x 10 min with PBS and mounted with Immu-Mount (9990402, Thermo Fisher Scientific).

### mRNA *in situ* hybridization

RNA in situ hybridization was performed using RNAScope Multiplex Fluorescent Reagent Kit v2 (323110, Advanced Cell Diagnostics) according to the manufacturer’s instructions. Small intestinal and colon tissues were collected, washed with PBS, and fixed in 4% paraformaldehyde for 24 h. The tissues were embedded in paraffin as Swiss rolls with a Leica Histocore Pearl tissue processing system. Then, 5 µm sections were cut using a microtome (Microm HM355, Thermo Fisher Scientific). Paraffin-embedded sections were deparaffinized in a xylene and ethanol series, and pre-treated with RNAScope Hydrogen Peroxide, Protease Plus, and Target Retrieval Reagents. Sections were hybridized with Lama5 (494911), Wif1 (412361), or Hhip (448441) probes (all from Advanced Cell Diagnostics). Signal amplification hybridization was performed, and signal detected with Opal 570 (FP1488001KT, Akoya Biosciences, 1:1000). Following hybridization steps, the tissue sections were immunostained. Samples were blocked with 5 % Normal Goat Serum in PBS for 45 minutes and incubated with primary antibodies [anti-E-Cadherin (610182, BD, 1:1000), anti-alpha smooth muscle Actin (ab5694, Abcam, 1:1000)] overnight at +4°C in blocking buffer. Samples were washed 3 x 10 minutes with PBS and incubated with secondary antibodies [Alexa Fluor 488 Goat anti-mouse IgG2a (115-547-186, Jackson ImmunoResearch, 1:500), Alexa Fluor 647 Goat anti-Rabbit IgG (A21245, Thermo Fisher Scientific, 1:1000)] and 1 µg/ml DAPI (D9542, Sigma-Aldrich). Samples were washed 3 x 10 minutes with PBS and mounted with Immu-Mount (9990402, Thermo Fisher Scientific).

### Gene regulatory networks, gene importance ranking and computational knockout simulations

To investigate the regulatory mechanisms underlying CAF differentiation, we employed the CellOracle pipeline [114] (version 0.20.0) to construct gene regulatory networks (GRNs) and perform *in silico* transcription factor (TF) knockout simulations. ScRNA-seq data was integrated with prior regulatory information to infer context-specific GRNs, which were then used to build linearized dynamical models approximating how gene expression responds to regulatory input. GRNs were constructed using the CellOracle Python package to model transcription factor (TF)–target gene interactions across fibroblast subtypes. We used previously described preprocessed scRNA-seq data, which included clustering and dimensionality reduction (UMAP) information, as well as raw expression counts, which were used for the modelling. For each fibroblast subtype, a context-specific GRN was inferred based on gene expression and a pre-defined base GRN. The context-specificity was achieved by trimming and refining the base GRN to fit it for the transcriptomic context for specific cell types or states. The trimming consisted of: (1) filtering the base GRN to retain TF-target pairs where both genes were expressed in the given cluster, (2) fitting a ridge-regression model to infer regulatory coefficients for each TF-target pair and (3) removing weak or statistically non-significant (p<0.001) edges based on statistical significance derived from the ridge-regression step. The gene importance ranking was directly performed from the trimmed, context-specific GRNs for each fibroblast subtype by calculating betweenness centrality scores to prioritize genes acting as major control points within the networks.

To allow computational knockout simulations, pseudotime was precomputed using scanpy’s diffusion pseudotime algorithm [115]. After pseudotime calculation, for each TF of interest, the expression value was individually set to 0.0 (knocked out), and the downstream effects on gene expression were estimated by approximating signal passing within the gene regulatory network. The model then calculates updated cell states (post-perturbation) by simulating the immediate transcriptional response (detailed in the original publication for CellOracle), and the updated cell states are then projected back onto the pre-existing low-dimensional cell manifold (e.g., UMAP used in this study), preserving biological interpretability.

### Human colorectal cancer study population

This study comprised a cohort of 315 patients with colorectal cancer (CRC) who underwent surgical treatment between 1998 and 2003 at the Department of Surgery, Helsinki University Hospital. Clinical data were retrospectively collected from patient records. Survival data were obtained through linkage with the Finnish Population Registration Centre and Statistics Finland. The median age at diagnosis was 67.8 years (interquartile range, 57.8–67.8 years). The median disease-specific survival was 5.9 years, with a range of 0.0–18.8 years. Ethical approval for the study was granted by the Surgical Ethics Committee of Helsinki University Hospital (Dnro HUS/1223/2021 and permission to use the tissue samples was granted by The Finnish Medicines Agency (Dnro FIMEA/2021/006901).

### Preparation of tumor tissue microarrays

Formalin-fixed, paraffin-embedded (FFPE) tumor samples from surgical specimens were retrieved from the archives of the Department of Pathology, University of Helsinki. All tumor samples were re-evaluated by an experienced pathologist using hematoxylin and eosin (H&E)-stained sections to confirm diagnoses and select representative tumor areas. Two punches, each 1.0 mm in diameter, were extracted from each tumor block using a semi-automated tissue microarray instrument (TMA) (Beecher Instruments, Silver Spring, MD, USA). Each TMA block was sectioned, resulting in two tissue cores per tumor sample.

### Tumor microarray immunohistochemistry

TMA blocks were sectioned at 4 µm thickness, mounted onto slides, and dried at 37°C for 12–24 hours. Antigen retrieval was performed using the Envision Flex target retrieval solution (pH 9, DM828, Agilent Dako, CA, USA) in a PreTreatment module (Agilent Dako) at 98°C for 15 minutes. Immunohistochemical staining was conducted with an Autostainer 480S (LabVision Corp., Fremont, CA, USA) utilizing the Dako REAL EnVision Detection System, Peroxidase/DAB+, Rabbit/Mouse. Sections were initially treated with Envision Flex peroxidase-blocking reagent (SM801, Dako) for 5 minutes to block endogenous peroxidase activity. Subsequently, the slides were incubated with anti-CTCHR1 antibody (ab256458, Abcam, Cambridge, United Kingdom), diluted 1:400 in Dako REAL Antibody Diluent (S2022, Dako), for 60 minutes. Afterward, slides underwent a 30-minute incubation with peroxidase-conjugated EnVision Flex/HRP rabbit/mouse reagent (SM802, Dako). Visualization was achieved using DAB chromogen (EnVision Flex DAB, DM827, Dako) for 10 minutes. Counterstaining was performed using Mayer’s hematoxylin (S3309, Dako). Normal colon served as the positive control.

### Interpretation of tumor microarray immunohistochemical staining

CTCHR1 immunoreactivity was independently assessed by two investigators (E.W.V. and J.H.), who were blinded to clinical outcomes. Immunostaining intensity was evaluated separately in tumor/epithelial cells and stromal cells, using a scoring system: negative (0), low (1), moderate (2), and strong immunoexpression (3). The highest score for each sample across all assessed compartments was considered representative of CTCHR1 expression. In cases of discrepancy between the two evaluators, consensus scoring was applied.

### Statistical analysis of tumor microarray staining

Survival analysis was done by the Kaplan-Meier method and the log-rank test. Disease-specific overall survival was counted from the day of surgery to the date of death from CRC, or until the end of follow-up. A p-value of 0.05 was considered significant. All tests were two-sided. All statistical analyses were done with SPSS version 29.0 (IBM SPSS Statistics version 29.0 for Mac; SPSS Inc., Chicago, IL, USA, an IBM Company).

### Immunofluorescent staining of human normal adjacent colon tissue

Normal adjacent colon tissue sections from the CRC study cohort described above were used for immunofluorescent staining. 5 µm-thick sections were deparaffinized in xylene and ethanol series. Antigen retrieval was performed using 1X Dako antigen retrieval solution pH 6 (S1699, Agilent) at +95°C for 20 min, followed by 30 min at room temperature (RT). Blocking was performed using TNB blocking buffer (SKU FP1012, Akoya Biosciences) for 30 min at RT. Primary antibodies anti-KCNN3 (APC-025, Alomone Labs, 1:100 dilution) and anti-PDGFRa (AF-307-NA, R&D Systems, 1:50 dilution) were incubated overnight at +4°C. Secondary antibodies Alexa Fluor 647 donkey anti-rabbit (A-31573, Thermo Fisher Scientific) and Alexa Fluor 594 donkey anti-goat (A11058, Thermo Fisher Scientific) were incubated in blocking buffer (1:500 dilution) for one hour at RT and washed 3×5 min in 0.1% TBS-Tween. The nuclei were stained with 10 µg/ml Hoechst 33324 (62249, Thermo Fisher Scientific) in PBS for 3 min. Slides were rinsed with TBS and water before mounting with Immu-Mount (9990402, Fisher Scientific).

### Imaging

Imaging of immunofluorescent human paraffin and mouse vibratome sections was done using the Zeiss Axio Imager M2 microscope using 20x magnification. Image analysis was conducted using Fiji ImageJ [116]. Images of immunofluorescent mouse paraffin sections, CRC TMA and RNA *in situ* hybridization sections were generated using 3DHISTECH Pannoramic 250 FLASH III digital slide scanner (20x and 40x magnification, respectively) at Finnish Genome Editing Center and analyzed with QuPath [117], CaseViewer (3DHISTECH) and CaseViewer, respectively.

### Gene set enrichment analysis

The GSEA was conducted using clusterProfiler [118] in R (version 4.2.3). Ranked lists for mouse MIF gene expression were created with the FindMarkers function in Seurat, comparing MIFs to all other clusters of the data using logfc.threshold = 0.1 and min.pct = 0.05. For humans, min pct was set to 0.25. For mouse GSEA, the mouse MSigDB ontology geneset M5 subcategory “molecular function (MF)” was used, and for human, the human MSigDB ontology geneset C5 subcategory “molecular function (MF)” was used [119–122]. Additionally, signatures of 250 top expressed genes were extracted from PDGFRA+ cells isolated from human colon muscle (PaC), ICCs isolated from human colon muscle (ICC), PDGFRA+ fibroblasts isolated from human colon mucosa (FIB), smooth muscle cells isolated from human colon muscle (SMC) [59] and enteric neurons (myenteric ganglia) isolated from human colon (EN) [58] and used for GSEA with a ranked list of human MIFs constructed with the FindMarkers function in Seurat comparing MIFs to all other clusters of the data using the default settings of the function.

### Trajectory and pseudotime analysis

Trajectory analysis was performed using the Monocle 3 algorithm [71]. Cells were clustered with resolution = 0.00012, which resulted in the best match with the Seurat clustering. Based on our analysis of normal human colon fibroblasts and the partitioning, where CAFs were in the same group as SAFs, MIFs, and SEMFs, we set SAF, MIF, and SEMF populations as originating clusters for the trajectory and pseudotime analyses performed using Monocle 3.

### CRC patient survival analysis

Survival analysis was performed using the Kaplan-Meier plotter website for colorectal cancer data (https://www.kmplot.com/analysis/index.php?p=service&cancer=colon) using the CAF subpopulation gene signatures (mean expression of selected genes). Autoselection for cutoff was used (percentile) for overall survival analysis. The cohorts included a total of 2.135 patients from the following datasets: GSE12945 (n = 62), GSE12294 (n = 155), GSE14333 (n = 123), GSE123985 (n = 91), GSE17538 (n = 232), GSE10800 (n = 53), GSE26682 (n = 331), GSE20540 (n = 35), GSE21595 (n = 37), GSE33114 (n = 90), GSE34489 (n = 33), GSE37892 (n = 65), GSE38832 (n = 70), GSE39582 (n = 514), GSE41258 (n = 185), GSE92921 (n = 59). Not all patients had the required clinical information for each analysis (e.g., CMS status, stage, etc.). In these cases, all patients with the required information were used in the analyses.

### Ethics statement

For scRNA-seq, mice were bred and maintained in the animal house facilities of BSRC Alexander Fleming under specific pathogen-free conditions, and experiments were performed in accordance with current European and national legislation (approved by BSRC Fleming’s Institutional Committee of Protocol Evaluation in conjunction with the Veterinary Service Management of the Hellenic Republic Prefecture of Attika). For validation experiments, mice were housed at the University of Helsinki Laboratory Animal Center according to national and international legislation and guidelines (license: KEK23-006, approved by the Research Ethics Committee on Animal Research). For the human CRC study cohort, ethical approval was granted by the Surgical Ethics Committee of Helsinki University Hospital (Dnro HUS/1223/2021 and permission to use the tissue samples was granted by The Finnish Medicines Agency (Dnro FIMEA/2021/006901).

## ACKNOWLEDGEMENTS

We thank Reeta Huhtala, Melissa Montrose, and Pia Saarinen for technical assistance, Finnish Genome Editing Center (supported by HiLIFE and the Faculty of Medicine, University of Helsinki, and Biocenter Finland) for scanning services, and Emilia Kuuluvainen for critical comments on the manuscript.

---

**S1 Fig. ScRNA-seq analysis of mouse colon fibroblasts.** (A) UMAP of the mouse colonic mesenchyme scRNA-seq performed in this study. (B) Feature plots depicting indicated marker genes for colon fibroblast subsets. (C) Violin plot showing unbiased marker genes for each identified fibroblast population. (d) Schematic representation of the datasets used for the integrated analysis of mouse mesenchymal cells. (E) Schematic representation of the datasets used for the integrated analysis of normal colon and CRC cells.

**S2 Fig. Characterization of identified mouse and human colon fibroblast subtypes.** (A) UMAPs split by individual datasets used for mouse integration analysis. (B) Dot plot depicting cell type marker genes in mouse colon. For fibroblast subsets, see Fig 1B. (C) UMAPs split by individual datasets used for human integration analysis. (D) Dot plot depicting cell type marker genes in human colon. For fibroblast subsets, see Fig 1D. (E) Feature plots showing a small *KIT*+ cluster representing ICCs (arrowhead) and a *CCL19*+ cluster representing reticular fibroblasts. (F) Dot plot of species-specific genes marking human but not mouse colon fibroblast subtypes (black genes in Fig 1D) in the mouse dataset. (G) Dot plot of species-specific genes marking mouse but not human colon fibroblast subtypes (black genes in S2H Fig) in the human dataset. (H) Dot plot depicting unbiased marker genes for each mouse fibroblast subpopulation (see Fig 1B for human genes). The markers shared between mouse and human are marked in red.

**S3 Fig. ScRNA-seq analysis of mouse small intestinal fibroblasts.** (A) UMAP of the mouse small intestinal fibroblasts and myocytes from scRNA-seq performed in this study. (B) Feature plot of the fibroblast marker *Pdgfra* and smooth-muscle marker *Myh11*. (C-E) Feature plots of indicated genes marking colon fibroblast subsets in the small intestinal mesenchyme dataset. C, markers for Top and Bottom SEMFs; D, markers for MAF subclusters; E, markers for SAFs.

**S4 Fig. Expression of mouse MAF and SEMF subset genes in human colon mesenchyme.** (A) Feature plot showing expression of mouse bottom MAF genes *TNC* and *CD55* in human data. (B) UPAM showing the difference in fibroblast subpopulations in right-sided (proximal) and left-sided (distal) colon stroma in human. (C) Feature plot of *WNT4* expression in human data. (D) Feature plots of mouse top SEMF marker expression in human. (E) Feature plots of mouse bottom SEMF marker expression in human.

**S5 Fig. Identification of CRC CAF subsets.** (A) UMAP showing contribution of individual datasets used for integration of normal adjacent and tumor tissue from CRC patients. (B) Dot plot depicting genes used for identifying non-mesenchymal cells in the integration of normal adjacent and tumor tissue. See also Fig 5D. (C) UMAP of the whole integrated CRC patient dataset (see Fig 5A) separated by sample origin normal adjacent tissue and CRC. (D) UMAPs visualizing the contribution of individual datasets used for the mesenchymal cell analysis of normal adjacent and tumor tissue (Fig 5B). (E) UMAPs of the individual samples included in the colon mesenchyme analysis (n = 53, > 50 cells and ≥ 5 clusters identified per sample) (F) Stacked bar plots depicting the stromal cell proportions per patient in normal adjacent colon and CRC. See also S11C Fig. (G) Feature plots showing a small *KIT*+ cluster representing ICCs, a *CCL19*+ cluster representing reticular fibroblasts, the antigen-presenting CAF marker CD74 enriching in the CCL19+ cluster, and proliferation marker MKI67 (arrowheads).

**S6 Fig. Marker genes of CRC CAF subsets and correlation of CAF subset signatures to survival.** (A-D) Dot plots of markers for (A) iCAF, (B) mCAF, (C) preCAF1 and (D) preCAF2 clusters. (E) Kaplan-Meier plot indicating correlation of the CAF subtype signature gene expression (see Fig 5E) with the probability of overall survival of CRC patients. mRNA-sequencing data derived from: GSE12945, GSE12294, GSE14333, GSE123985, GSE17538, GSE10800, GSE26682, GSE20540, GSE21595, GSE33114, GSE34489, GSE37892, GSE38832, GSE39582, GSE41258, GSE92921.

**S7 Fig. Presence of CAFs and normal colon fibroblasts in CRC and normal tissue.** (A) Projections of CAF subtype signatures (Fig 6A) onto normal human colon spatial transcriptomics data. (B) Dotplot depicting specific marker gene signatures for normal colon fibroblast subsets. Only markers not expressed in CAFs were selected for this analysis. (C) Projections of normal-specific fibroblast subtype signatures from B onto normal colon tissue and CRC tumor tissue spatial transcriptomics data. (D) Gene regulatory network assisted gene importance ranking. Transcription factors marked in red/purple. (E) Monocle3 clustering of the integrated human mesenchymal scRNA-seq data. Clusters 3, 5, 9 and 10 represent Seurat clusters iCAF, preCAF2, preCAF1 and mCAF, respectively. (F) Monocle3 partitioning analysis. CAFs partitioned together with clusters 6, 8 and 14 representing the Seurat clusters SAF, MIF and SEMF, respectively, indicating a shared trajectory between CAFs and these normal-like fibroblasts. (G) Trajectory and (H) pseudotime analyses of the fibroblast clusters. (I) mRNA expression signature of potential transcription factors *PRRX1*, *MAFB* and *TWIST1* driving CAF identity identified in Oracle analysis (see Fig 5B).

**S8 Fig. Representation of the CAF subsets across all CRC samples.** (A) UMAPs showing CAF subpopulations in each CRC patient. Red boxes indicate the samples excluded from subsequent analysis (<50 CAFs).

**S9 Fig. Correlation of CAF subtype representation with molecular and clinical characteristics of CRC patients.** (A) Stacked bar plot of CAF subtype proportions across all 40 samples included in the CAF analysis. (B-C) Stacked bar plots of CAF proportions in different CMS subtypes. B, overall; C, per patient. (D-F) Enrichment of specific CAF subtypes in CMS1-4. D, preCAF2s in CMS1s; E, preCAF1s in CMS3; F, mCAFs in CMS4. (G-H) Stacked barplot of CAF subtype proportions in iCMS2 and iCMS3. G, average across patients; H, per patient. (I-J) Stacked barplot of CAF subtype proportions in left or right-sided tumors. I. overall; J, per patient. (K) Stacked barplot of CAF subtype proportions in proximal colon, distal colon and rectum. (L) Enrichment of preCAF1 in the rectum. (M-N) Stacked bar plots of CAF subtype proportions in microsatellite stable (MSS) and microsatellite instable (MSI-high) CRCs. M, overall; N, per patient. (O) Enrichment of preCAF2 in MSI-H patients. (P-Q) Stacked barplot of CAF subtype proportions in different tumor stages. P, overall; Q, per patient. (R) Enrichment of iCAFs proportion early stage (stage 1-2) as compared to late stage (3/4) CRCs. A p-value < 0.05 was considered statistically significant. All enrichments of CAF subtypes with p < 0.1 are shown.

**S10 Fig. Correlation of CAF subtype representation with genetic characteristics of CRC patients.** (A) Stacked barplot of CAF subtype proportions in *APC* wild-type and mutant CRC in all patients (left) and per patient (right). (B) Stacked barplot of CAF subtype proportions in *KRAS* wild-type and mutant CRC in all patients (left) and per patient (right). (C) Boxplot of mCAF proportion in cancers with or without *KRAS* mutation. (D) Stacked barplot of CAF subtypes in *KRAS* wild-type and mutant patients separated stage. (E) Boxplot of mCAFs abundance in early and late-stage CRC with or without *KRAS* mutation. (F) Stacked barplot of CAF subtype proportions in *TP53* wild-type and mutant CRC in all patients (left) and per patient (right). (G) Boxplot of preCAF1 proportion in cancers with and without *TP53* mutation. (H) Stacked barplot of CAF subtype proportions in *BRAF* wild-type and mutant CRC in all patients (left) and per patient (right). (I) Stacked barplot of CAF subtype proportions in *PIK3CA* wild-type and mutant CRC in all patients (left) and per patient (right). (J) Boxplot of preCAF1 proportion in cancers with or without *PIK3CA* mutation. A p-value < 0.05 was considered statistically significant. All enrichments of CAF subtypes with p < 0.1 are shown in this plot.

**S11 Fig. CTHRC1 is a marker for CRC CAFs.** (A) Immunohistochemical staining of normal adjacent tissue from CRC patients. Note absence of CTHRC1 staining in normal colon stroma but weak expression in lymphocyte-like cells in normal colon mucosa in two out of four patients (arrowheads). (B) Correlation of stromal CTHRC1 staining density to survival of CRC patients. (C) Mesenchymal cell type proportions in normal colon tissue and in early and late-stage CRC. CAFs marker with red box. (D-E) Expression of *CTHRC1* (D) and *FAP* (E) mRNA across CRC CMSs (Sidra-LUMC AC-ICAM dataset). Note highest expression levels in CMS4 patients. Statistical comparisons of normalized mRNA expression levels were performed using Student’s t-tests, followed by Bonferroni adjustment for multiple testing. (G-I) Kaplan-Meier plots depicting the correlation between mRNA expression of the panCAF signature (*CTHRC1, INHBA, THBS2, WNT2, FAP*), *CTHRC1* and *FAP* with overall survival in early (stage 1-2) and late-stage (3-4) CRC. (J-L) Kaplan-Meier plots showing correlation of panCAF signature (*CTHRC1, INHBA, THBS2, WNT2, FAP*), or *CTHRC1* and *FAP* alone with CRC patient survival across CRC CMSs. mRNA-sequencing data derived from: GSE12945, GSE12294, GSE14333, GSE123985, GSE17538, GSE10800, GSE26682, GSE20540, GSE21595, GSE33114, GSE34489, GSE37892, GSE38832, GSE39582, GSE41258, GSE92921.

**S1 Table. Mouse scRNA-seq datasets used for integration.** Details of the scRNA-seq datasets used for the mouse integration analysis, including filtering parameters and the number of analyzed cells for each dataset.

**S2 Table. Human colon fibroblast and CRC CAF signatures.** Transcriptional signatures of each human fibroblast and CRC CAF subset used for spatial transcriptomics and signature UMAPs.

**S3 Table. Summary of published mouse and human colon fibroblast subsets.** Detailed presentation of the previously published colon fibroblast subsets, including original annotations, markers, localization, references, and corresponding annotations provided in this study.

**S4 Table. Summary of published CRC CAF subsets.** Detailed presentation of the previously published CRC CAF subsets, including original annotations, markers, localization, references, and corresponding annotations provided in this study.

