## Supplemental Figure 1 for "High-resolution integrative analysis allows characterization and spatial annotation of normal and cancer-associated colon fibroblasts"

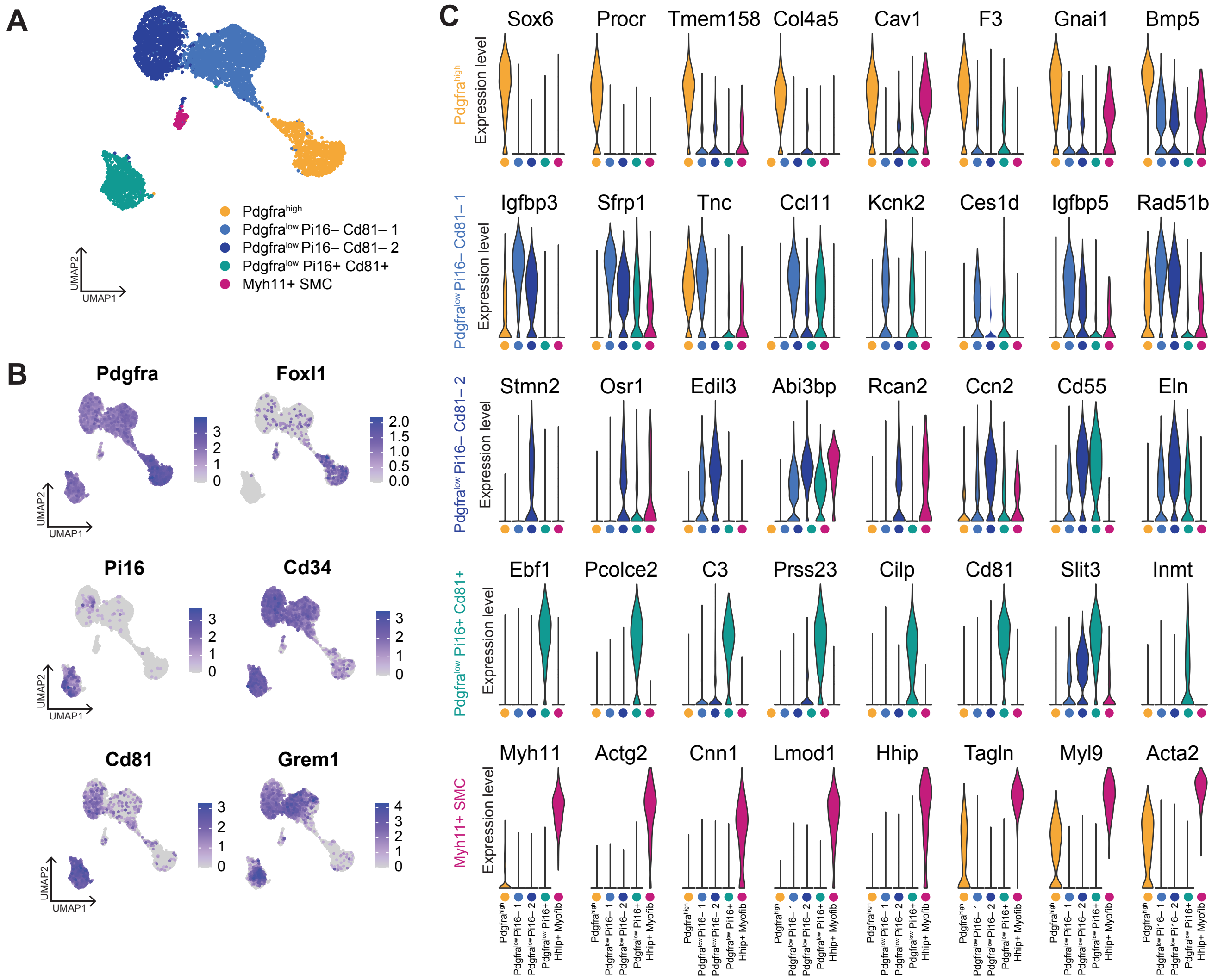


D

| Dataset | Collected tissue | Enrichment strategy |
|---|---|---|
| Roulis | colon mesenchyme | – |
| Jasso | colon mesenchyme | EPCAM- CD45- TERT119- |
| Kinchen | colon mesenchyme | EPCAM- CD45- |
| Brügger | colon | – |
| This study | colon mesenchyme | EPCAM- CD45- |

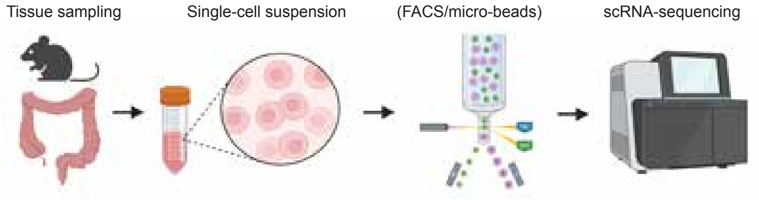


E

| Dataset | Collected tissue | Reagent kit |
|---|---|---|
| CRC16 | normal colon & CRC | Chromium Single Cell 3′ v3 |
| KUL3 | normal colon & CRC | Chromium Single Cell 3′ v2 |
| KUL5 | normal colon & CRC | Chromium Single Cell 5′ |
| SMC | normal colon & CRC | Chromium Single Cell 3′ v2 |
| JSC | CRC | Chromium Single Cell 5′ |

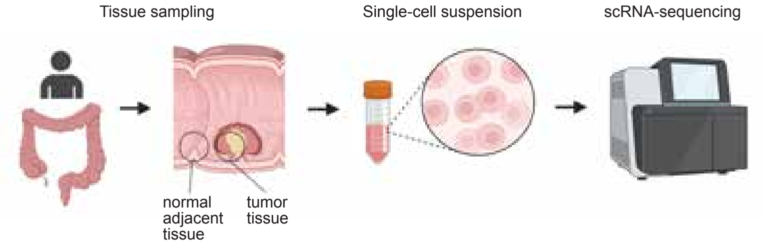
