## Supplementary figures and images for "High-resolution integrative analysis allows characterization and spatial annotation of normal and cancer-associated colon fibroblasts"

### Supplemental Figure 2

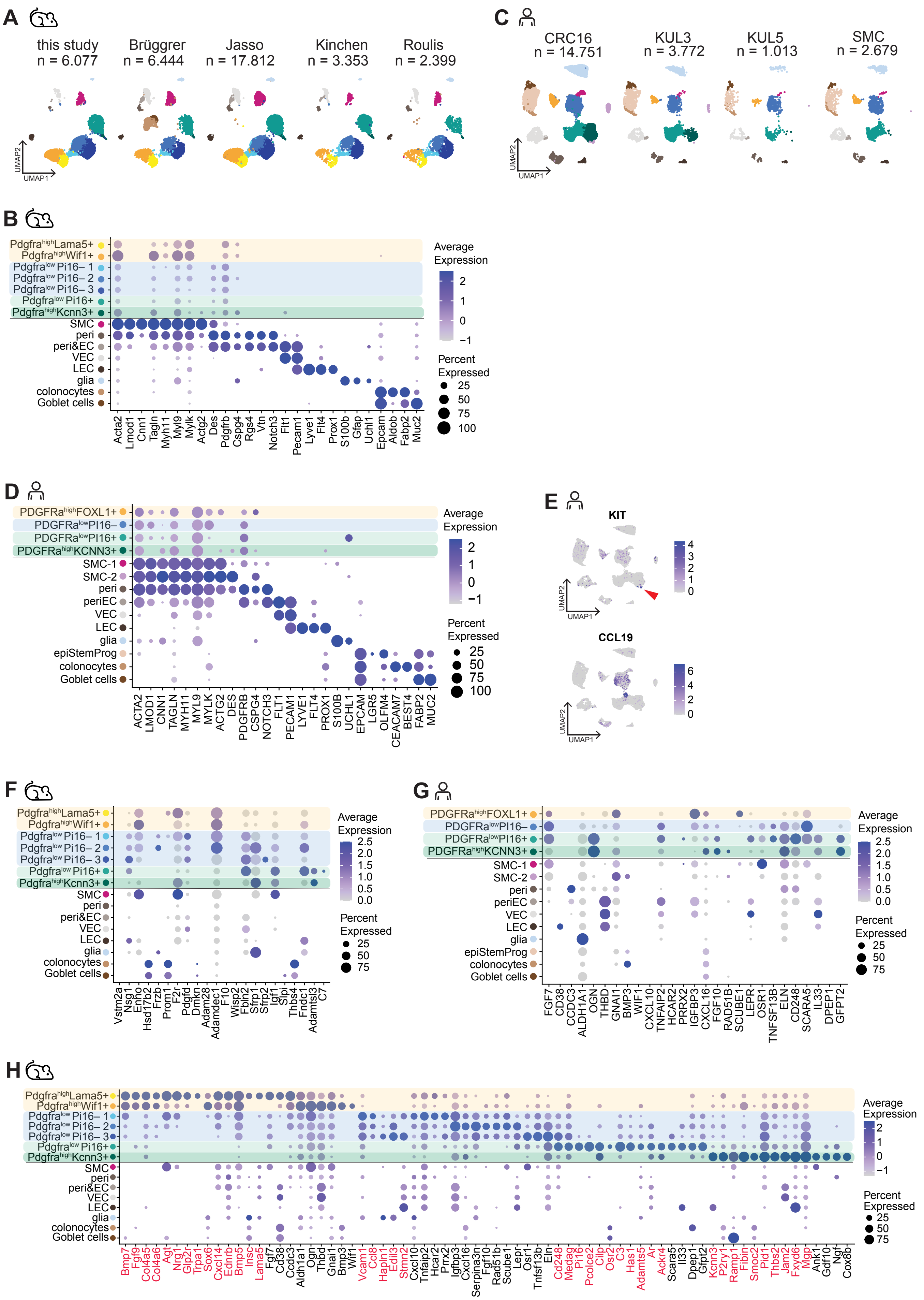

### Supplemental Figure 3

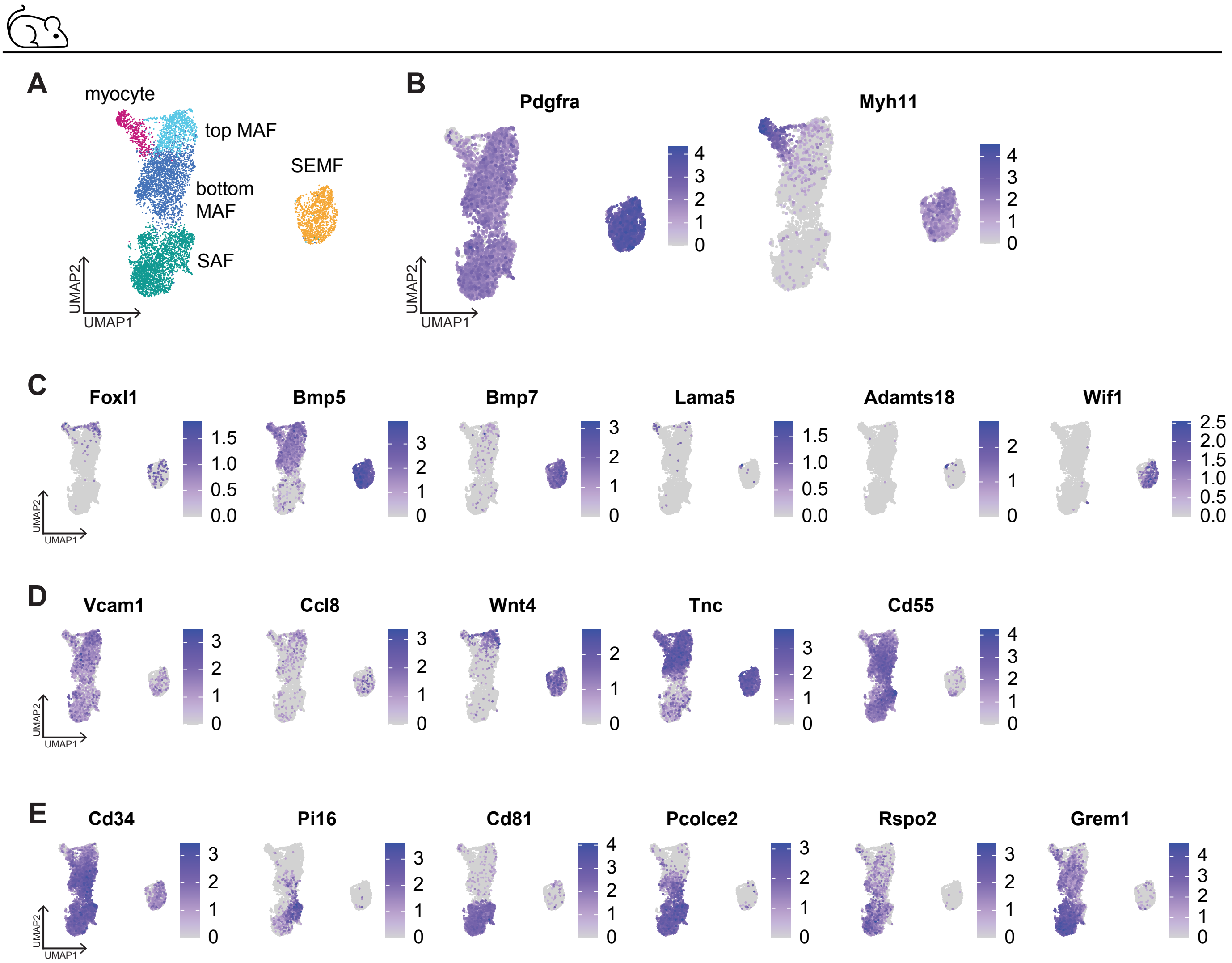

### Supplemental Figure 4

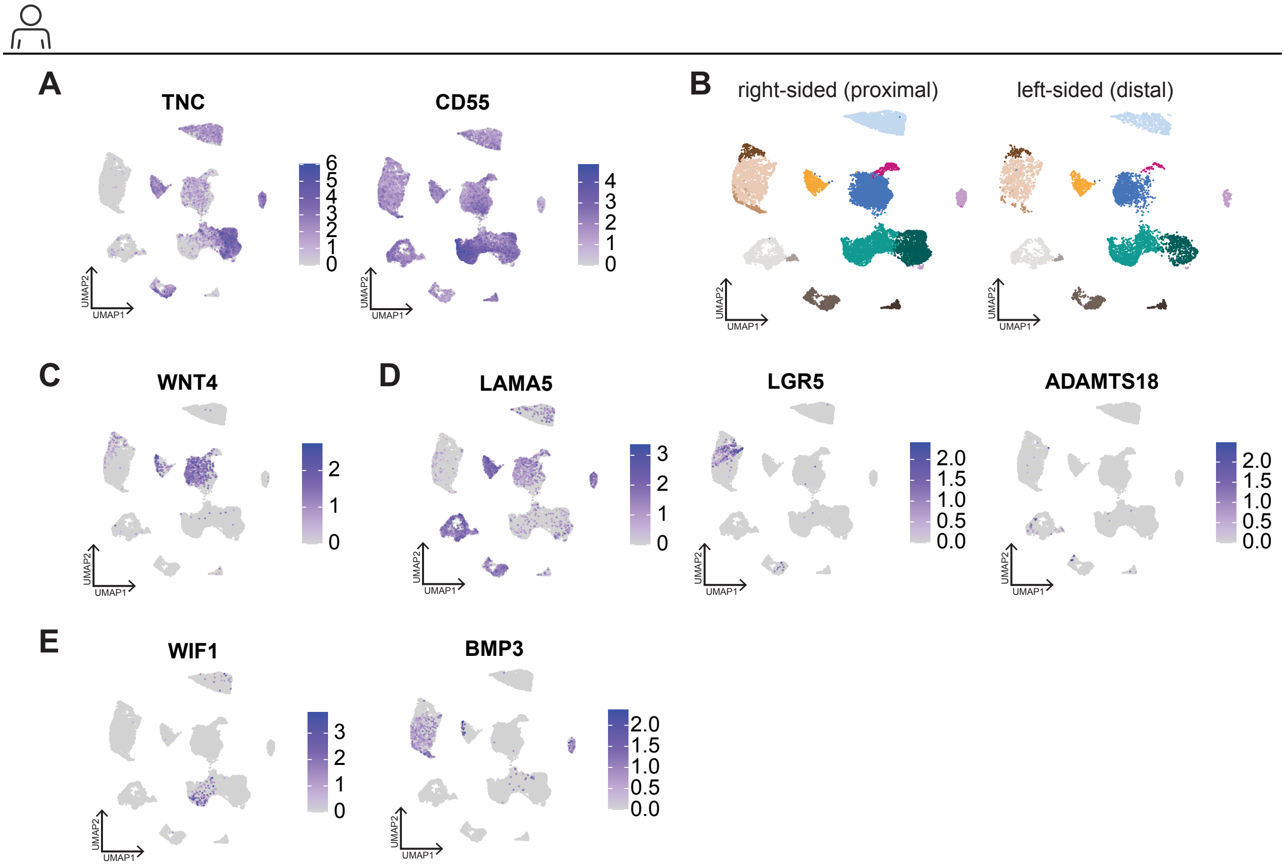

### Supplemental Figure 5

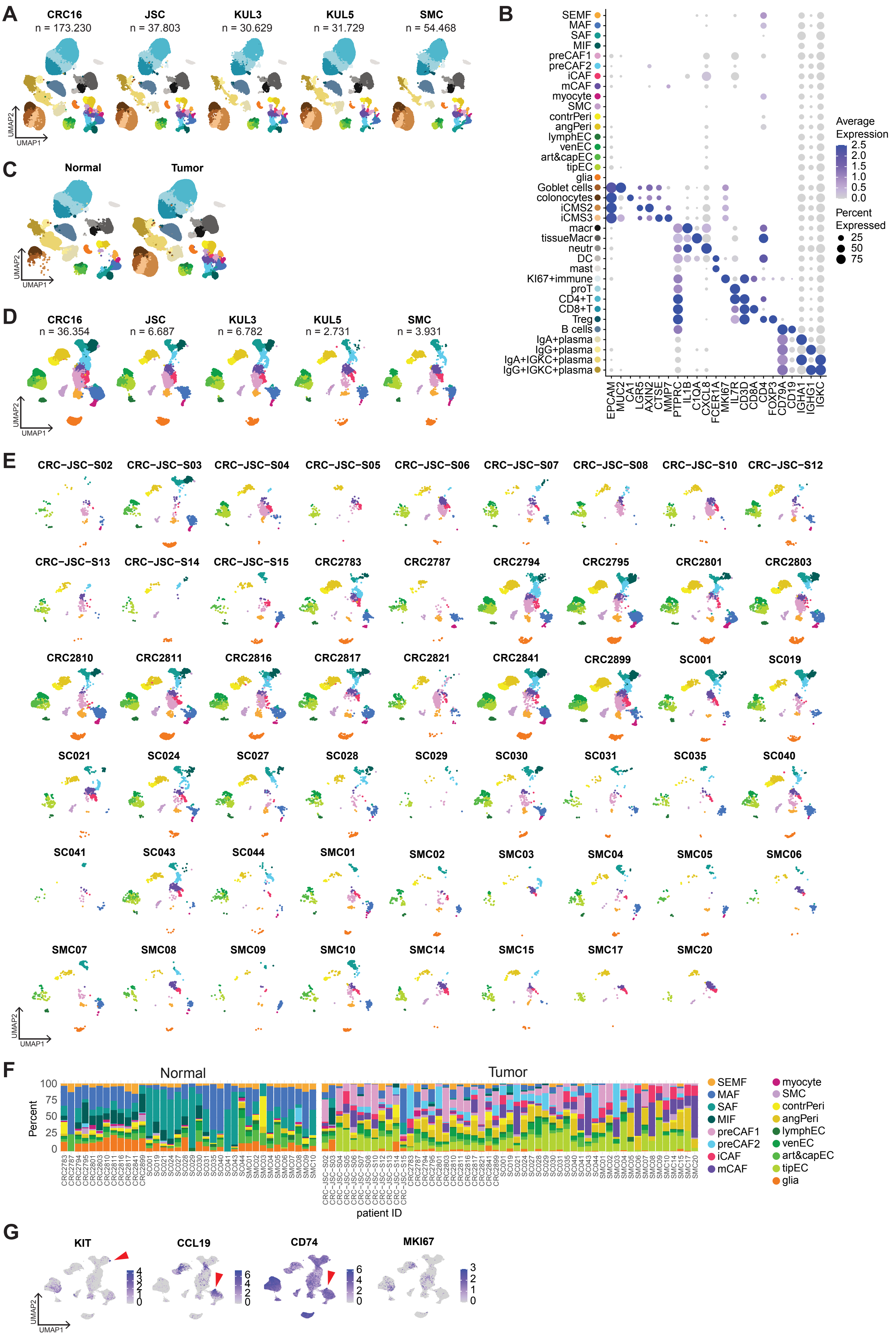

### Supplemental Figure 6

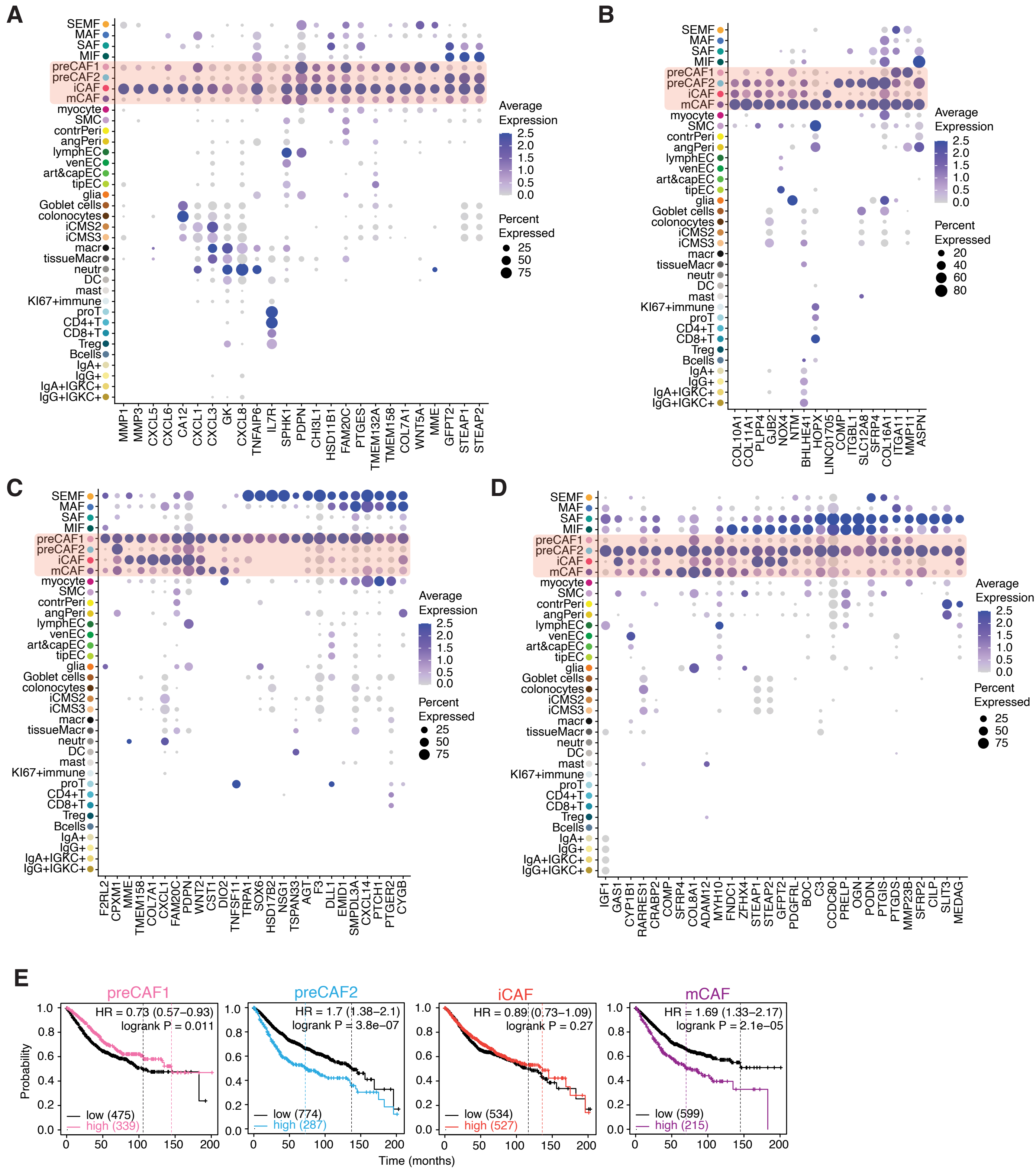

### Supplemental Figure 7

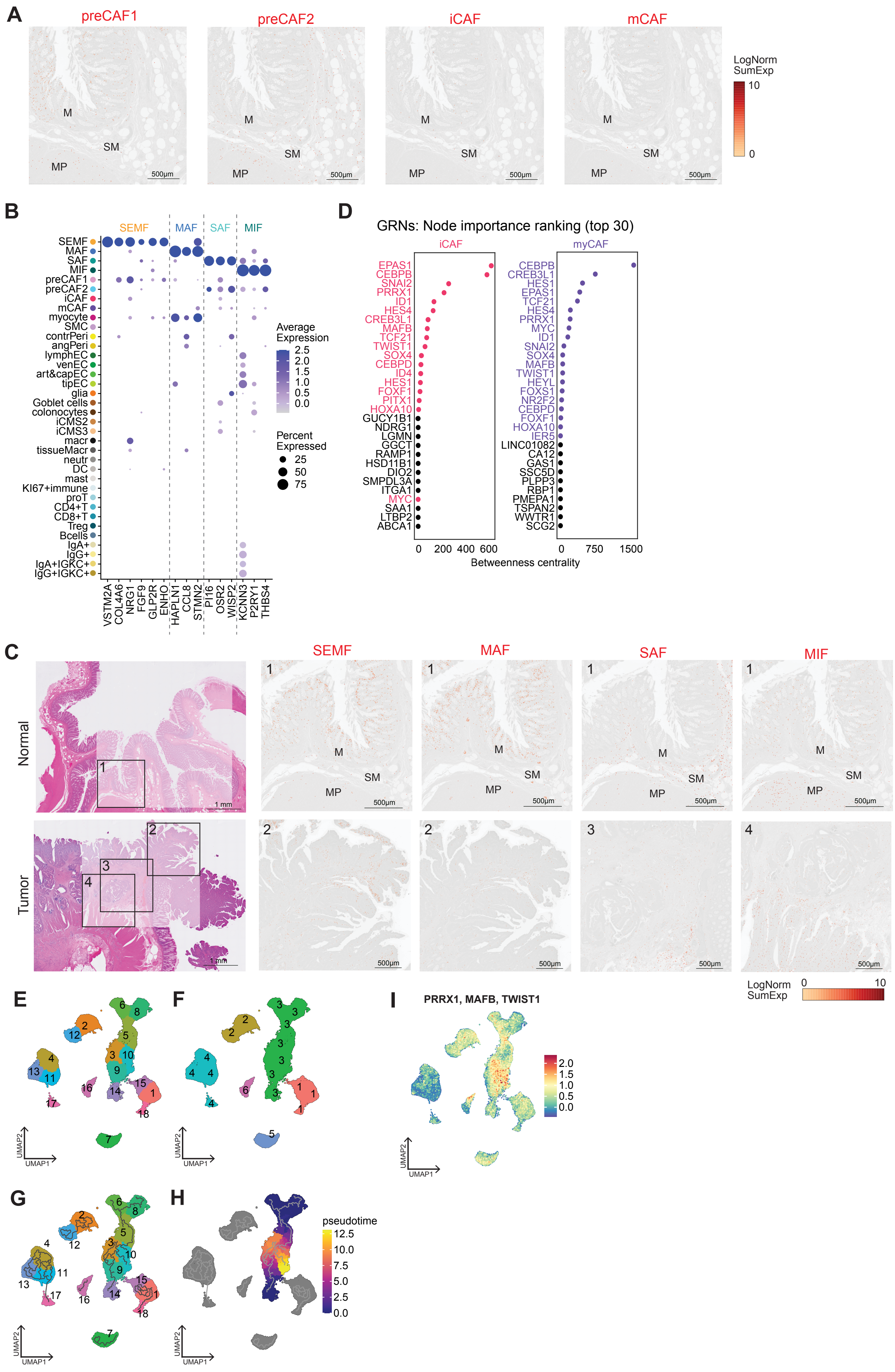

### Supplemental Figure 8

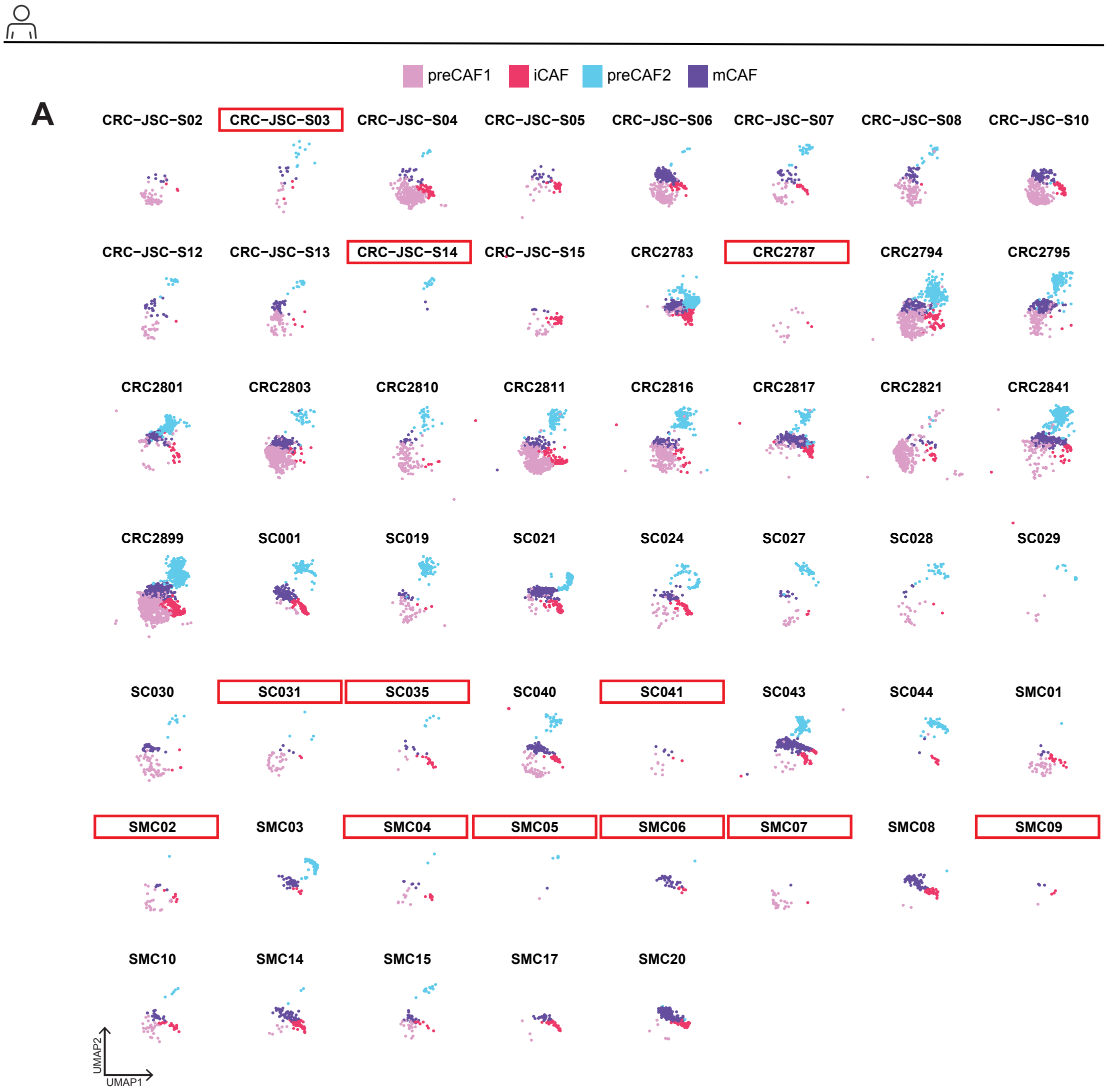

### Supplemental Figure 9

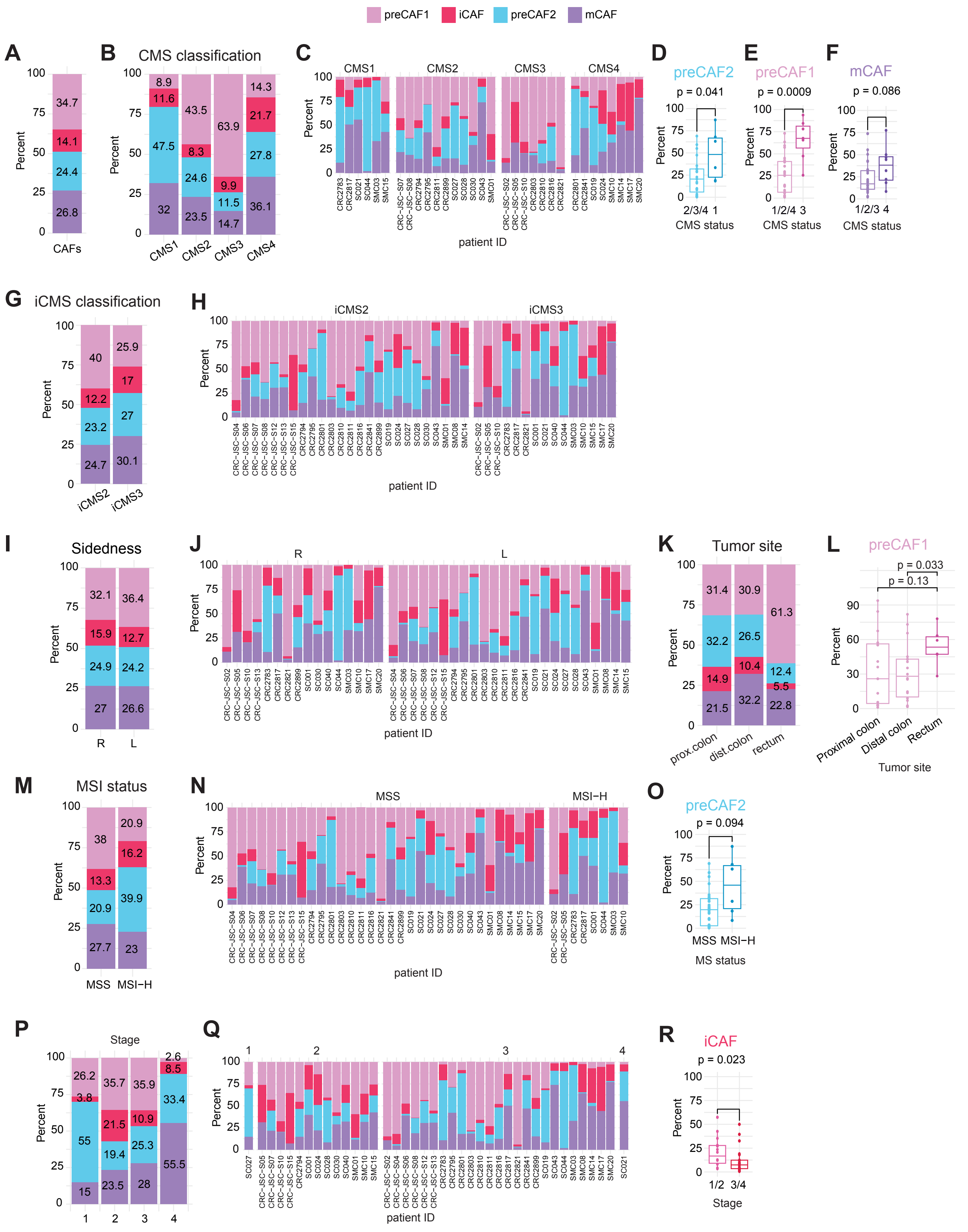

### Supplemental Figure 10

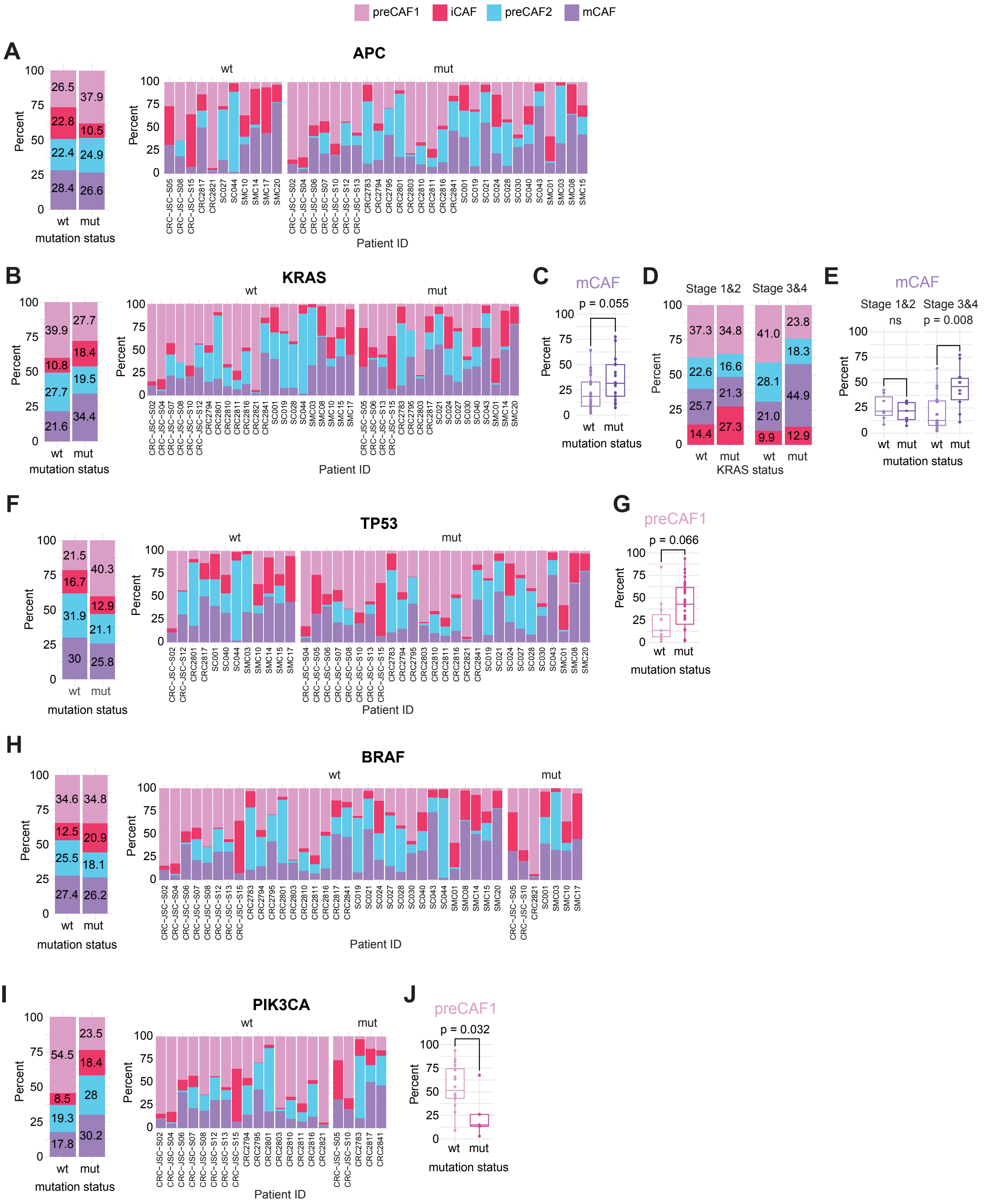

### Supplemental Figure 11

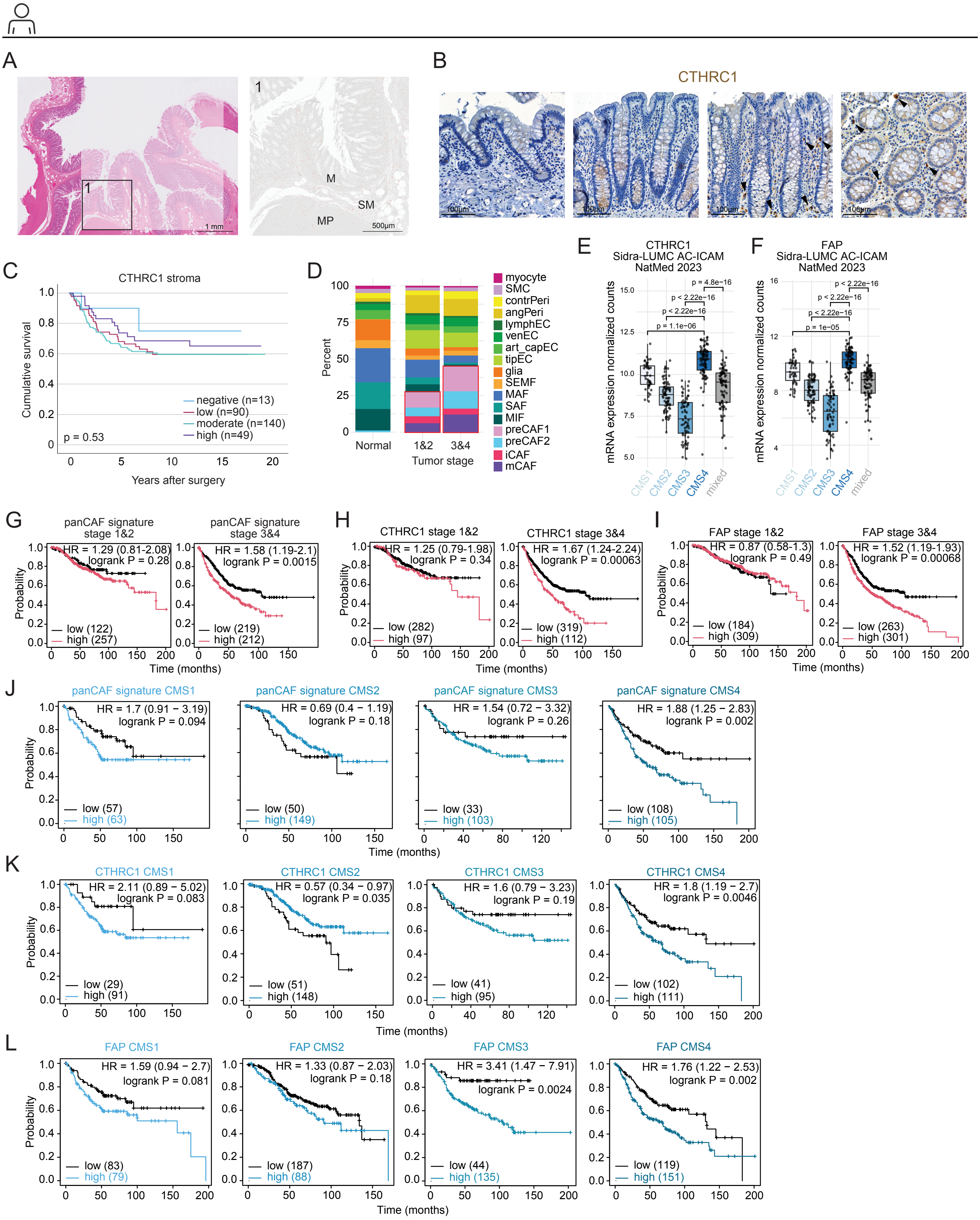
